# Spatial proteomic mapping of human nuclear bodies reveals new functional insights into RNA regulation

**DOI:** 10.1101/2024.07.03.601239

**Authors:** Boris J.A. Dyakov, Simon Kobelke, B. Raktan Ahmed, Mingkun Wu, Jonathan F. Roth, Vesal Kasmaeifar, Zhen-Yuan Lin, Ji-Young Youn, Caroline Thivierge, Kieran R. Campbell, Thomas F. Duchaine, Benjamin J. Blencowe, Archa H. Fox, Anne-Claude Gingras

**Affiliations:** Lunenfeld-Tanenbaum Research Institute, Mount Sinai Hospital, Sinai Health, Toronto, ON M5G 1X5, Canada; Department of Molecular Genetics, University of Toronto, Toronto, ON M5S 1A8, Canada; School of Human Sciences, The University of Western Australia, Crawley, WA 6009, Australia; Donnelly Centre, University of Toronto, Toronto, ON M5S 3E1, Canada; Program in Molecular Medicine, The Hospital for Sick Children, Toronto, Ontario M5G 0A4, Canada; Rosalind and Morris Goodman Cancer Institute, McGill University, Montreal, QC H3A 1A3, Canada; Department of Biochemistry, McGill University, Montreal, QC H3G 1Y6, Canada; Department of Statistical Sciences, University of Toronto, Toronto, ON M5G 1Z5, Canada; Department of Computer Science, University of Toronto, Toronto, ON M5S 2E4, Canada; Ontario Institute of Cancer Research, Toronto, ON M5G 0A3, Canada; Vector Institute, Toronto, ON M5G 1M1, Canada

**Author notes:** Correspondence (A.H.F.), (A.-C.G.).

**Keywords:** BioID, proximity-based labelling, mass spectrometry, nuclear bodies, paraspeckle, nuclear speckle, nucleolus, Cajal body, PML body, QKI, NEAT1, splicing

## Abstract

Nuclear bodies are diverse membraneless suborganelles with emerging links to development and disease. Explaining their structure, function, regulation, and implications in human health will require understanding their protein composition; however, isolating nuclear bodies for proteomic analysis remains challenging. We present the first comprehensive proximity proteomics-based map of nuclear bodies, featuring 140 bait proteins (encoded by 119 genes) and 1,816 unique prey proteins. We identified 641 potential nuclear body components, including 131 paraspeckle proteins and 147 nuclear speckle proteins. After validating 31 novel paraspeckle and nuclear speckle components, we discovered regulatory functions for the poorly characterised nuclear speckle- and RNA export-associated proteins PAXBP1, PPIL4, and C19ORF47, and revealed that QKI regulates paraspeckle size. This work provides a systematic framework of nuclear body composition in live cells that will accelerate future research into their organisation and roles in human health and disease.

## Introduction

Nuclear bodies are membraneless structures with roles in gene regulation and nuclear and genome organisation^1–5^. Formed mainly *via* phase separation, they influence cellular processes by concentrating specific factors (proteins and RNAs)^6,7^ that regulate many essential processes related to gene expression^8–12^. Accordingly, nuclear bodies and their components play roles in development and disease. For example, the PML body is organised by PML, a protein frequently translocated in promyelocytic leukaemia and implicated in the antiviral response^13,14^, the Cajal body component SMN is mutated in spinal muscular atrophy^15^ and paraspeckles are essential for early mouse development^16^ and are implicated in neurodegenerative disease^17^. However, leveraging these roles to treat human diseases will require a deeper understanding of their composition.

While imaging-based colocalisation studies have revealed markers of each structure^18–20^, the approach is not amenable to unbiased discovery. Cell lysis and biochemical fractionation coupled with mass spectrometry (MS) has helped define the compositions of some nuclear bodies, notably the nucleolus^21^ and the nuclear speckle^22^. However, this requires purifying intact structures, which is challenging for membraneless organelles.

Proximity-based labelling (such as BioID and APEX^23^) has recently emerged as a means of characterising the protein compositions of specific organelles^24–26^, including membraneless organelles like the centrosome^27^ and cytoplasmic RNA bodies and granules^28^. We previously used APEX-MS/-sequencing to describe the protein and RNA composition of nuclear bodies^29^ and an updated atlas of nuclear speckles was recently generated using tyramide signal amplification (TSA)-MS^30^. However, systematic profiling of nuclear bodies using multiple ‘baits’—which is critical to distinguish whether identified ‘preys’ are true components of a protein structure^31^—has not been performed.

In this study, we used BioID and mass spectrometry to generate the first systematic inventory of human nuclear body components, which can be used to gain functional insights into these enigmatic structures.

## Results

### A spatial map of human nuclear bodies

Since existing Gene Ontology (GO) terms for some nuclear bodies are incomplete (*e.g.,* paraspeckles) or lack full validation (*e.g.,* nuclear speckles), we reviewed the existing literature, including all prior microscopy-based^18–20^ and proteomic^22,30^ surveys of nuclear body composition, to compile curated lists of known nuclear body proteins (**Fig. 1a, Supplementary Table 1**). We tagged at least 15 baits for each well-characterised nuclear body (nucleoli, nuclear speckles, PML bodies, Cajal bodies, and paraspeckles), the nuclear pore/envelope, and chromatin or RNA-associated proteins with the BioID enzyme BirA*^32^. We also chose baits associated with Sam68 nuclear bodies, histone locus bodies, and perinucleolar compartments (classified under “RNA-associated” baits).

**Fig. 1.**
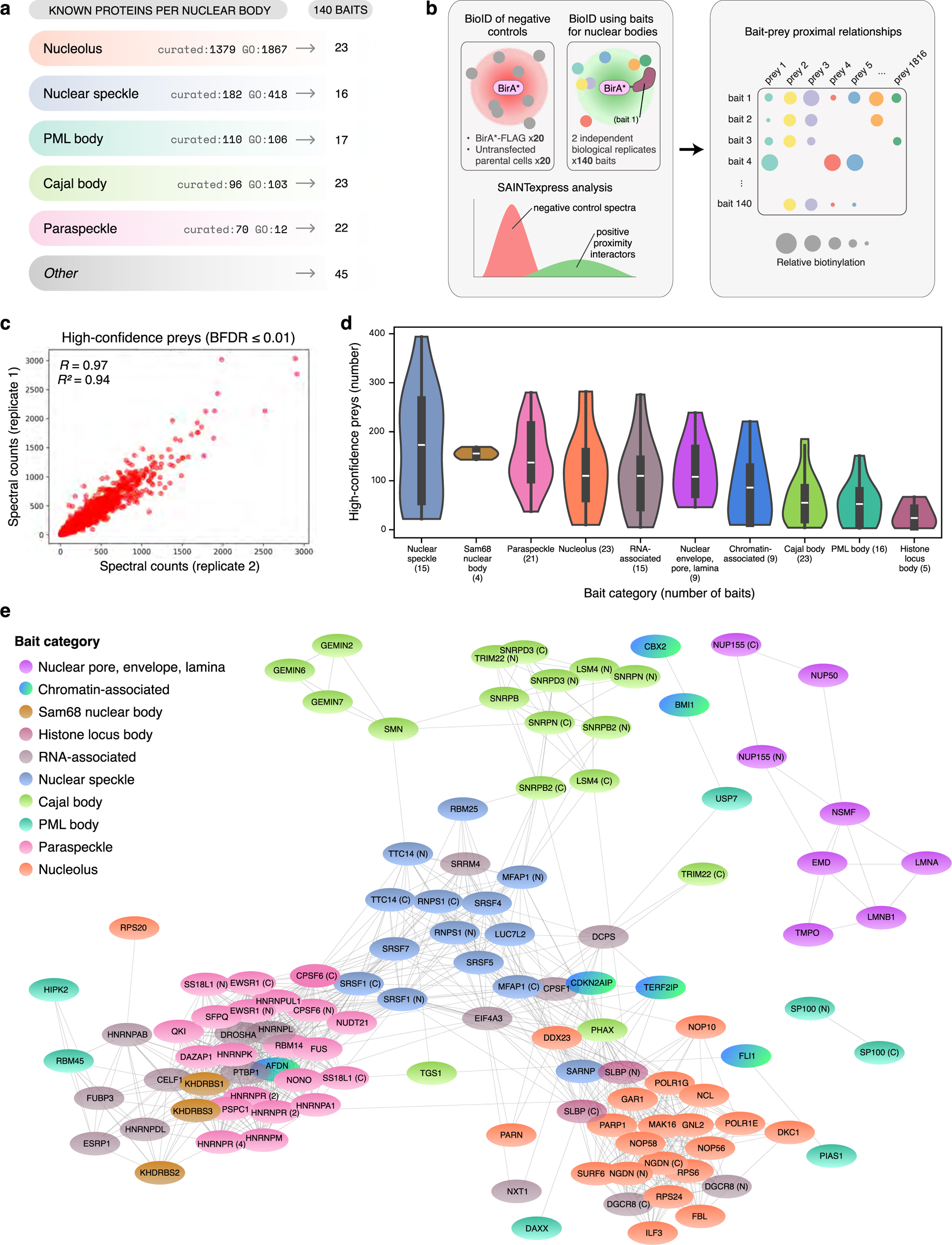
A BioID map of diverse nuclear bodies. **a**, Overview of the baits used to profile various human nuclear bodies. **b**, The BioID experimental workflow. **c**, Scatter plot of the Pearson correlations between the spectral counts of each high-confidence prey in two independent biological replicates. **d**, Violin plots of the distributions of high-confidence preys recovered by baits in various categories. **e**, Network of bait-bait Pearson correlations of unfiltered (pre-SAINTexpress) prey spectral counts. The colours indicate bait categories. Correlations > 0.5 are shown.

In total, HEK293 Flp-In T-REx inducible stable cell pools expressing 146 baits (representing 123 unique genes; **Fig. 1a, Supplementary Table 2**) were subjected to two independent biological replicates of BioID. We included negative controls with each batch of samples (untransfected parental cells and pools expressing BirA*-FLAG; *n*=20 each), enabling us to filter high-confidence preys from proteins that were non-specifically or endogenously biotinylated or non-specifically bound to the affinity matrix using SAINTexpress^33^ (**Fig. 1b**). Six baits with poor prey recovery were excluded from further analysis. The remaining 140 (representing 119 genes) recovered 14,877 proximal associations with 1,816 high confidence preys (*i.e.,* those with a Bayesian false discovery rate (BFDR) ≤ 0.01; **Supplementary Table 3 and the <u>ProHits-web online resource</u>**). For quality control purposes, approximately half of the baits were analysed by fluorescence microscopy after biotin labelling to confirm their expression, nuclear localisation, expected nuclear body staining pattern, and biotinylation (**Extended Data Fig. 1a**).

The spectral counts for preys recovered in both biological replicates of their baits were highly correlated (*R*^2^ = 0.90 and 0.94 for the unfiltered and high-confidence spectral counts, respectively; **Fig. 1c, Extended Data Fig. 1b**). The number of high-confidence preys varied between baits (range: 3–394 preys, mean: 106, median: 89; **Extended Data Fig. 2a**) and structures (**Fig. 1d**). The baits selected for each nuclear body type revealed significant enrichment in relevant terms (**Extended Data** Fig. 2b**, Supplementary Table 4**), had highly correlated prey recovery profiles (**Fig. 1e, Extended Data Fig. 1c**), and were usually reciprocal proximal interactors (**Extended Data Figs. 3 and 4**), indicating that the dataset provides a robust survey of nuclear body composition in HEK293 cells.

### Non-negative matrix factorisation defines the composition of various nuclear bodies

We used non-negative matrix factorisation (NMF) to assign high-confidence preys to nuclear bodies and other spatially or functionally associated clusters^24,28^. NMF factorises spectral count data into ‘bait’ and ‘prey’ matrices defined by a specified number (*k)* of clusters. We selected *k*=19 as the optimal number of clusters (based on precision-recall analysis and manual curation, see Methods) and set minimum thresholds for primary cluster assignment (≥ 5% of the highest score in the cluster), and secondary or tertiary cluster assignment (≥ 70% of the highest score across all clusters; see Methods; **Extended Data Fig. 5a–c;** full NMF results in **Supplementary Table 5**).

In a network generated using pairwise Pearson correlations between the NMF scores of all preys, annotations based on primary NMF assignments resulted in distinct clusters (**Fig. 2a**), which were classified based on their GO terms by gene set enrichment analysis (GSEA; **Fig. 2b, c**). Related NMF clusters were positively correlated in their NMF scores (**Extended Data Fig. 5d**), terms were enriched across multiple clusters, and numerous preys were assigned to secondary or tertiary clusters (**Extended Data Fig. 5e**), consistent with our existing knowledge of nuclear body composition, function, and relative organisation inside the cell. For example, the cluster for nuclear speckles, which are thought to support splicing of transcripts from relatively highly expressed genes^10,12^, contained many splicing-related proteins. Accordingly, clusters 6 (splicing) and 12 (nuclear speckle) were highly enriched for the same splicing-related GO terms (**Fig. 2c**), positively correlated (*R* = 0.64, the highest between any two clusters; **Extended Data Fig. 5d**), and closely clustered (**Fig. 2a**). Consistent with this, 31 proteins in cluster 12 were also assigned to cluster 6 (**Extended Data Fig. 5e**).

**Fig. 2.**
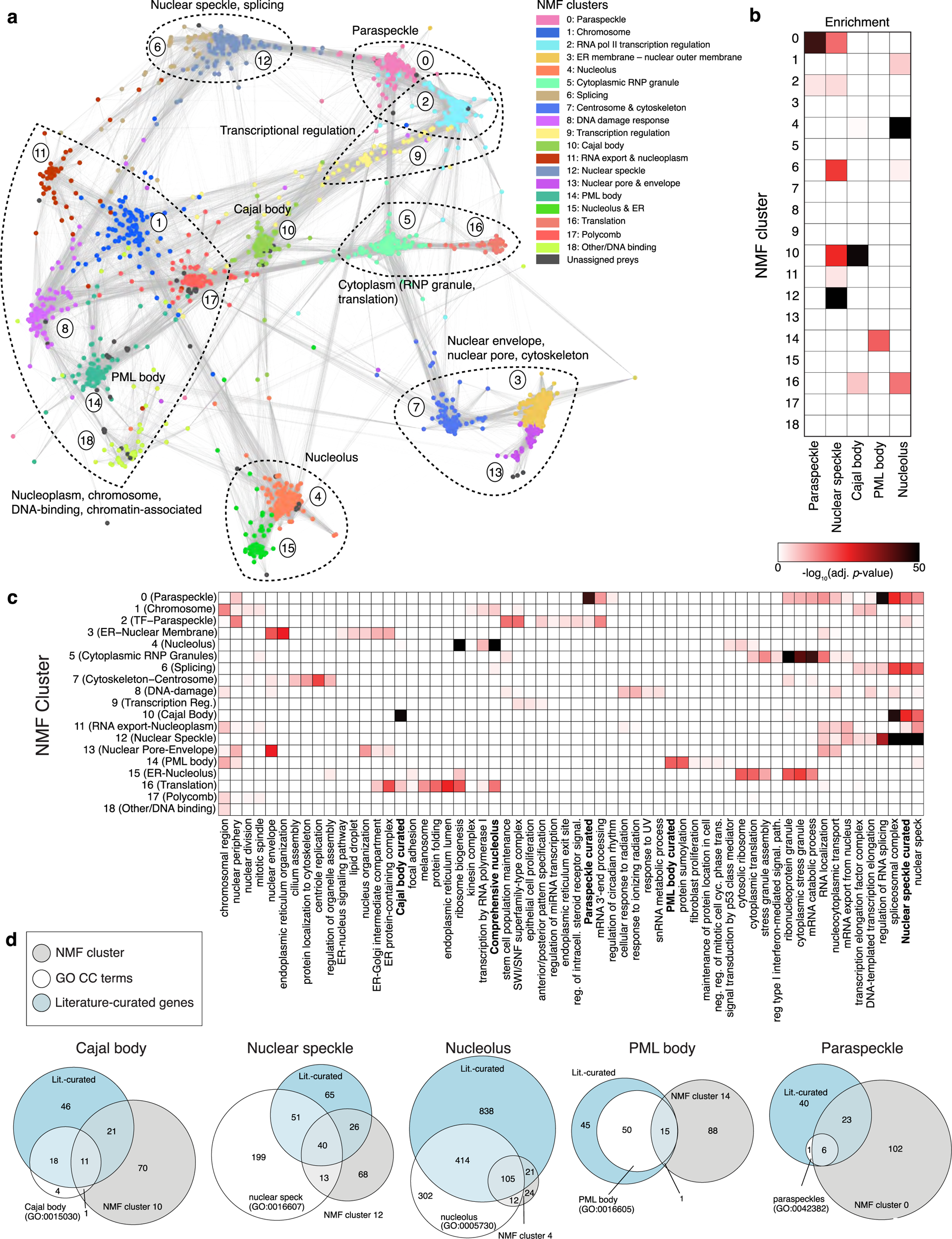
Non-negative matrix factorisation assigns preys to specific nuclear bodies. **a**, Network of Pearson correlations ≥ 0.6 (denoted by grey edges) between the NMF scores of all high- confidence preys. The colours indicate primary NMF cluster assignments. **b**, Summarized Gene Set Enrichment Analysis (GSEA) results for each NMF cluster against curated terms for the nuclear bodies listed. **c**, GSEA results for each NMF cluster. Columns indicating enrichment in custom literature-curated terms for specific nuclear bodies are indicated in bold. Full results are provided in Supplementary Table 4. **d**, Venn diagrams showing the overlap between known and novel human nuclear body proteins in their NMF clusters.

Paraspeckles (cluster 0), which include many transcriptional regulators^34^, were closely associated with general regulators of mRNA transcription (cluster 2). Cluster 2 contained a distinct set of proteins from cluster 9, which also contained transcriptional regulators, indicating that we can spatially resolve distinct groups of proteins with similar roles. Paraspeckles also contain many RNA-binding proteins and are often adjacent to nuclear speckles^35,36^; this is indicated by the edges (representing prey-prey correlations) between clusters 0 and 12 (**Fig. 2a**), and their common GO term enrichments (**Fig. 2c**).

We identified large clusters representing the Cajal body (102 proteins), nuclear speckle (147 proteins), nucleolus (162 proteins), PML body (103 proteins), and paraspeckle (131 proteins). This included high proportions of proteins that are known nuclear body components (**Supplementary Table 1**) or are linked to associated GO terms, with particularly strong performance for the Cajal body, nuclear speckle, and paraspeckle clusters (recovering 33%, 36%, and 41% of known proteins, respectively; **Fig. 2d**). Interestingly, aside from the nucleolar clusters, most proteins in each cluster were not previously annotated as components of their assigned nuclear body, including 46% (68/147) of nuclear speckle, 71% (70/103) of Cajal body, 78% (102/131) of paraspeckle, and 85% (88/103) of PML body proteins (vs 15% (24/162) of nucleolar proteins). These results also suggest that the nucleolus may be undersampled in our study, given its large size and high number of known proteins.

The top 50 paraspeckle proteins (by NMF score) include well-known markers (FUS, RBM14, TARDBP, *a.k.a.* TDP-43, and SFPQ) and functionally consistent novel components, including RNA-binding proteins and transcriptional regulators (**Fig. 3a**), while the top 50 nuclear speckle proteins included various splicing factors (**Fig. 3b**). Top Cajal body proteins included key markers (COIL, SMN1), spliceosomal snRNPs, and SMN complex proteins (**Extended Data Fig. 6a**), consistent with its roles in snRNP biogenesis, maturation, and recycling^11^. Top PML body proteins included key markers (PML, DAXX) and many proteins involved in the small ubiquitin-related modifier (SUMO) pathway, consistent with the implicated role of PML bodies in regulating SUMOylation^13^ (**Extended Data Fig. 6b**). All top 50 proteins in the nucleolus cluster were known nucleolar proteins (**Extended Data Fig. 6c**).

**Fig. 3.**
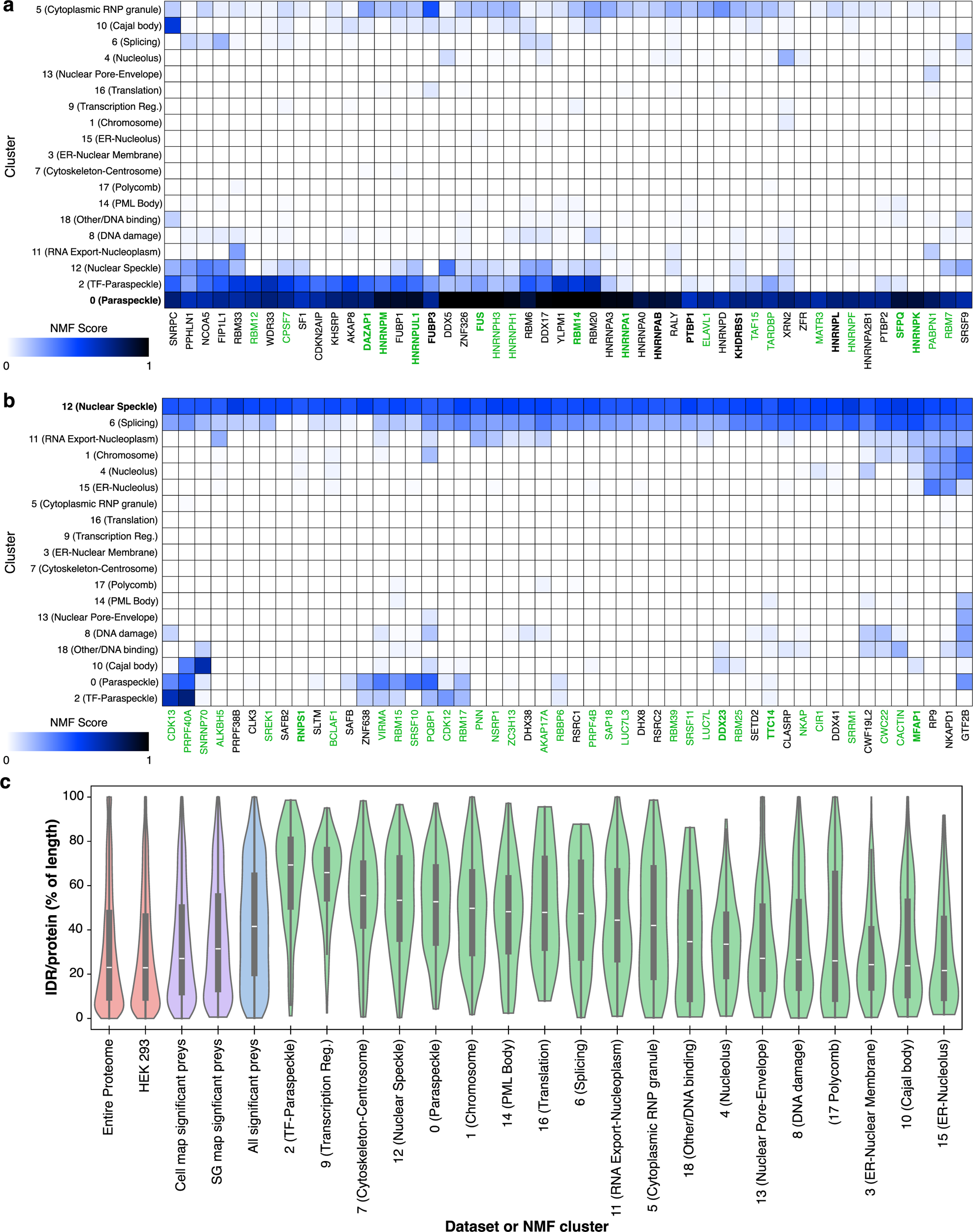
Proteins in NMF clusters are associated with specific nuclear bodies and enriched in IDRs. **a**, A hierarchical clustering of the NMF scores of the top 50 genes in the paraspeckle cluster (0). Known paraspeckle proteins are in green, and baits in this study are indicated in bold. **b**, A hierarchical clustering of the NMF scores of the top 50 genes in the nuclear speckle cluster (12). Known nuclear speckle proteins are in green, and baits in this study are indicated in bold. **c**, Violin plot of the relative proportions of the proteins in each NMF cluster with IDRs, arranged by descending median percent disorder. Their proportions in the entire human proteome, the HEK293 proteome, all high-confidence preys (BFDR ≤ 0.01) in this dataset, and all high-confidence preys from two other BioID datasets (human cell map (BFDR ≤ 0.01)) and the stress granule/P body map (AvgP ≥ 0.95)) are also shown. Pairwise statistically significant differences in IDR distributions between the clusters (by Wilcoxon-Mann-Whitney test) are provided in Supplementary Table 5.

Protein domain enrichment analysis of each NMF cluster further supports our ability to spatially map distinct cellular locations, with enrichment in domains functionally related to each nuclear body or process (*e.g.* RNA recognition motifs in paraspeckles and nuclear speckles; DEAD and helicase domains in the nucleolus and nuclear speckles; **Extended Data** Fig. 7a **and Supplementary Table 5**). This analysis expands our cluster annotation by enabling more detailed protein classification. For example, clusters 2 and 9 are enriched with transcriptional regulators (**Supplementary Table 5**) and are highly correlated (**Extended Data Fig. 5d**), while cluster 2 is also highly correlated with the paraspeckle cluster. While cluster 9 is enriched in proteins containing C2H2 zinc fingers, cluster 2 is enriched with bromodomain-, homeodomain-, and PAS domain-containing proteins, supporting the notion that paraspeckles regulate gene expression in specific developmental stages and tissue types, or under dynamic stress-responsive conditions^37^.

Liquid-liquid phase separation occurs through multivalent protein-protein and protein-nucleic acid interactions, often mediated by intrinsically disordered regions (IDRs) in the proteins. Therefore, we compared the predicted percentages of IDRs in each protein in our dataset against those in the human proteome, in other BioID datasets^24,28^, and in each cluster (**Fig. 3c**). The transcriptional regulation-associated clusters (2 and 9) had the highest percentage of IDRs, highlighting the importance of IDRs and phase separation in gene regulation^38,39^. Except for the Cajal body cluster, all nuclear body clusters had significantly higher IDR percentages compared to that predicted for the HEK293 proteome (adjusted (adj.) *p <* 0.0005, Mann-Whitney U test). LLPhyScore, a machine learning-based model for predicting phase separation based on physical interactions and features^40^, determined that all nuclear body clusters contained proteins predicted to phase separate. The paraspeckle cluster showed the highest predicted propensity to phase separate, with the median for the entire cluster in the top 10% of proteins in the entire proteome (**Extended Data Fig. 7b**). Altogether, our dataset greatly expands the known nuclear body proteome, providing numerous novel components and literature-supported inter-cluster connections and domain enrichments, including phase separation-related features.

### Novel paraspeckle and nuclear speckle proteins colocalise and interact with their predicted nuclear bodies

Since the nuclear speckle and paraspeckle had the best depths of coverage (high recovery of known proteins, numerous novel components; **Fig. 2d**), and high correlations between baits chosen for each structure (**Fig. 1e, Extended Data Figs. 1c and 3**), we focused our validation and further characterization efforts on them. We transiently expressed enhanced green fluorescent protein (EGFP)- tagged candidate paraspeckle (24) and nuclear speckle (9) proteins in HEK293 and/or HeLa cells, and assessed their colocalisation with the essential paraspeckle scaffolding RNA NEAT1^41^ and the nuclear speckle marker SRRM2^42,43^. NONO was used as a positive control for paraspeckles and SRSF7 for nuclear speckles; NONO was also used as an additional paraspeckle marker in 6/10 HeLa experiments. As the transcription inhibitor actinomycin D induces larger, more spherical nuclear speckles that facilitate imaging of their cores^44^, we included this treatment for approximately half of our nuclear speckle candidates. In total, 20/24 paraspeckle candidates colocalised with NEAT1 in HeLa and/or HEK293 cells. All nine nuclear speckle candidates colocalised with SRRM2 (representative images in **Fig. 4a, b, Extended Data Fig. 8a, b**, and **Supplementary Fig. 1**).

**Fig. 4.**
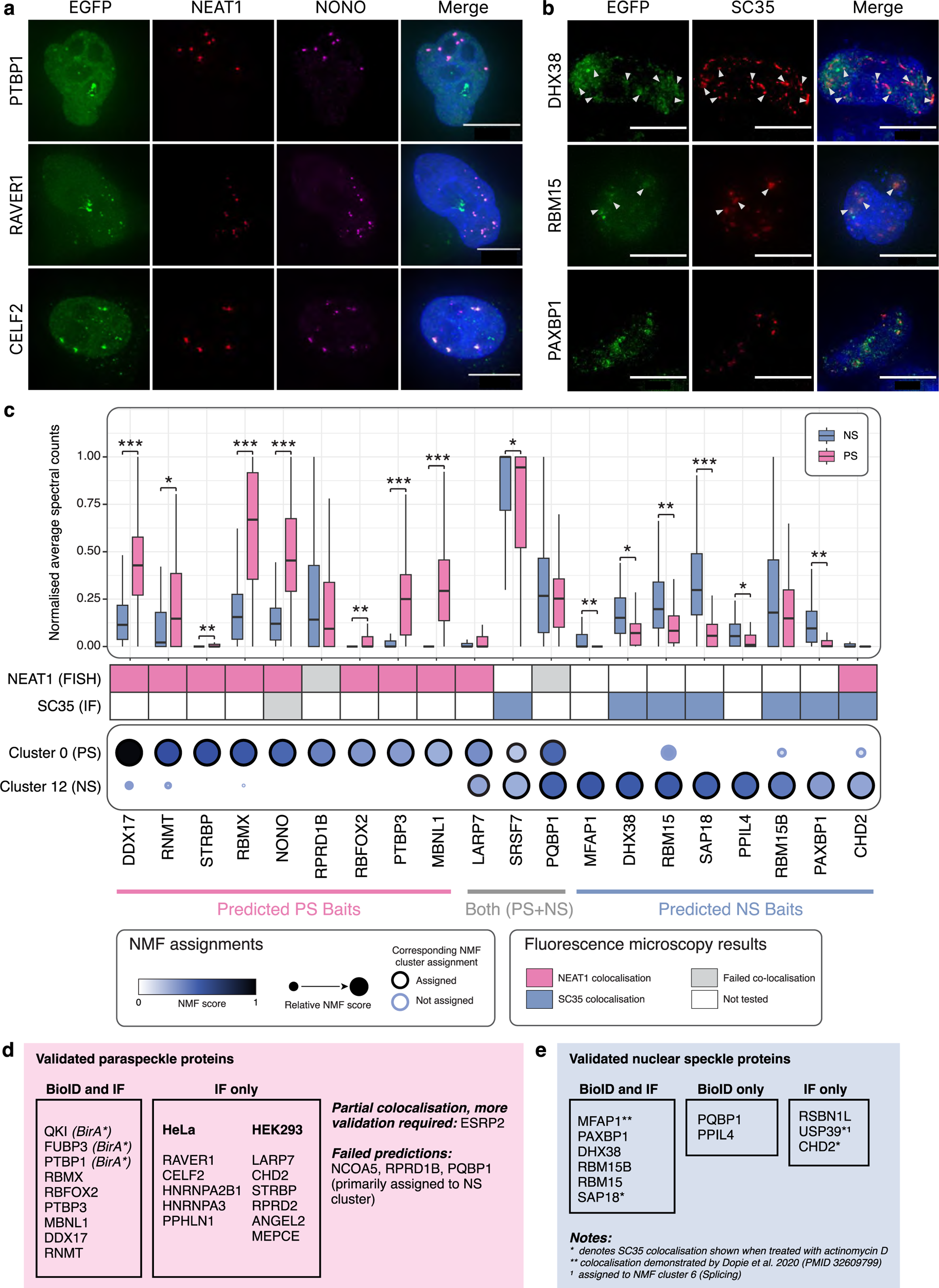
Validation of predicted paraspeckle and nuclear speckle proteins. **a**, Representative fluorescence images of validated paraspeckle proteins in HeLa cells. The merged images include DAPI to visualize the nucleus. Scale bars = 10 µm. **b**, Representative fluorescence images of nuclear speckle proteins validated in HEK293 cells. The merged images include DAPI staining to visualize the nucleus. White arrowheads indicate colocalised foci. Scale bars = 10 µm. **c**, Paraspeckle and nuclear speckle candidate validation. Significant differences in the per-bait distributions of recovered paraspeckle and nuclear speckle proteins are indicated as: *, *p <* 5×10^-2^, **, *p <* 5×10^-8^, ***, *p <* 5×10^-15^ by Wilcoxon signed-rank test. **d**, Summary of paraspeckle protein validation. **e**, Summary of nuclear speckle protein validation.

We also tagged 20 previously-unknown paraspeckle and nuclear speckle proteins with miniTurbo^45^ for reciprocal BioID (**Supplementary Tables 6 and 7**). Though many baits recovered known and NMF cluster-defined preys linked to both nuclear bodies, predicted paraspeckle and nuclear speckle proteins recovered significantly more spectra (**Fig. 4c**) and unique preys (**Extended Data Fig. 8c**) from their predicted nuclear body.

Reciprocal BioID supported the validation of 14 proteins already shown to colocalise with paraspeckle or nuclear speckle markers, as well as two additional nuclear speckle-associated proteins (PQBP1 and PPIL4; **Fig. 4d, e**, **Supplementary Table 8**). We validated several top-scoring proteins in the paraspeckle cluster, including PPHLN1, FUBP3, DDX17, HNRNPA3, PTBP1, and HNRNPA2B1 (**Fig. 3a**). FUBP3, PTBP1, and QKI were also used as BirA*-tagged baits in the main dataset and correlated strongly with other paraspeckle baits (**Fig. 1e, Extended Data Fig. 1c**). Similarly, MFAP1 was among the top 50 nuclear speckle proteins by NMF score (**Fig. 3b**) and was also used as a BirA*- tagged bait where it was highly correlated with nuclear speckle baits (**Fig. 1e, Extended Data Fig. 1c**). During the course of this study, MFAP1 was independently identified and validated as a nuclear speckle protein by TSA-MS^30^; the results of this study have been included in our curated list (**Supplementary Table 1**).

Notably, NCOA5, one of top 50 paraspeckle proteins, did not colocalise with NEAT1 in HeLa cells, nor did the paraspeckle candidates RPRD1B and PQBP1. However, all three proteins formed foci in cells, suggesting the existence of paraspeckle sub-assemblies lacking NEAT1, an interpretation supported by recent evidence showing that paraspeckles can assemble on a non-NEAT1 lncRNA in early mouse embryos^46^ (**Supplementary Fig. 1a, d**). Reciprocal BioID of RPRD1B and PQBP1 recovered paraspeckle proteins. Notably, PQBP1 was primarily assigned to the nuclear speckle cluster (**Fig. 4c** **and Extended Data Fig. 8c**), further highlighting the complex relationship between these nuclear bodies.

Overall, our > 87% validation rate for predicted paraspeckle and nuclear speckle proteins (29/33) demonstrates that proximity labelling and clustering analyses is effective for the identification of new nuclear body components.

### Novel nuclear speckle-associated proteins regulate transcription and splicing

Nuclear speckles associate with sites of active transcription and contain splicing, transcription, and export-related components^30,47^. Our nuclear speckle baits recovered several largely uncharacterised proteins, including three that we further analysed. PPIL4 (a peptidylprolyl isomerase; these enzymes participate in signalling, transcription, pre-mRNA processing, and mRNA decay^48^) and PAXBP1 (which promotes histone 3 lysine 4 methylation, a marker of active transcription^49,50^) clustered with nuclear speckle proteins. C19ORF47, an uncharacterized protein with no known functional domains (**Extended Data Fig. 9a**) that was recently reported to localise within nuclear speckles^51^, was the second highest-ranked protein in cluster 11 (RNA export and nucleoplasm; **Supplementary Table 5**). Reciprocal BioID with these candidates recovered proteins involved in mRNA transcription, splicing and processing, and export, including many known or NMF-predicted nuclear speckle proteins (**Fig. 5a**, also see **Fig. 4c** and **Extended Data Fig. 8c**).

**Fig. 5.**
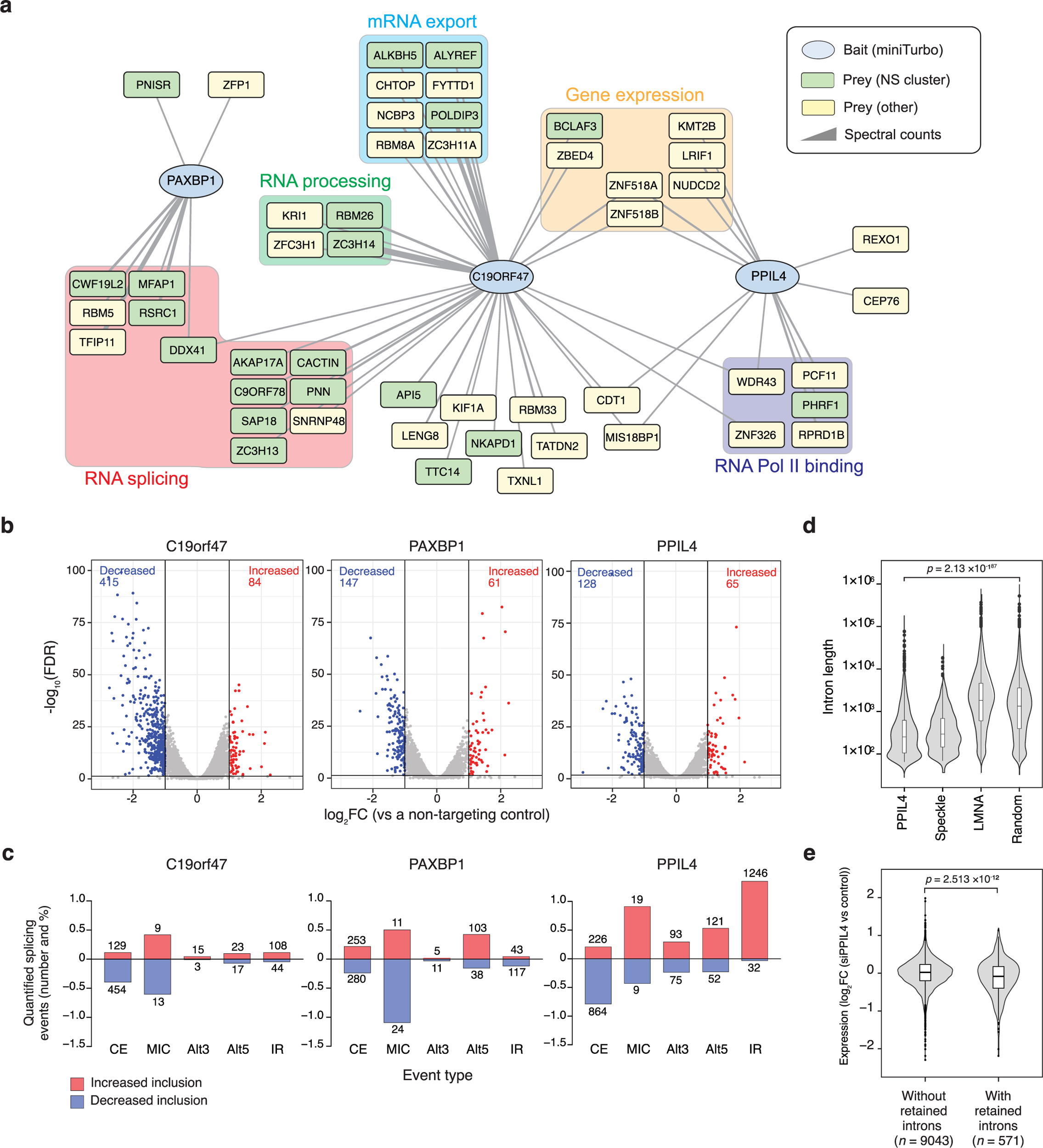
Nuclear speckle proteins regulate gene expression and alternative splicing. **a**, Network showing the high-confidence proximal preys (BFDR ≤ 0.01) of PAXBP1, PPIL4, and C19ORF47. **b**, Differential expression analysis after C19ORF47, PAXBP1, and PPIL4 depletion compared to a non-targeting control. **c**, Percentage and number of types of alternative splicing events showing at least 10% increased or decreased inclusion following C19ORF47, PAXBP1, or PPIL4 depletion (CE, cassette exon; MIC, microexon; Alt5/3, alternative 5′ or 3′ splice site; IR, intron retention). **d**, Lengths of PPIL4-retained introns compared to nuclear speckle-/lamina-associated introns from Barutcu, Wu *et al.* 2022^29^ or randomly selected introns. **e**, Relative expression changes in genes with and without PPIL4-retained introns following PPIL4 knockdown.

To examine their possible effects on transcriptional regulation and mRNA splicing, we depleted each with short interfering (si)RNA pools (**Extended Data Fig. 9b, c**) and performed poly(A)+- selected RNA-Seq. Depleting each candidate resulted in broad and significant changes in mRNA levels compared to non-targeting siRNA-transfected controls (**Fig. 5b**), particularly for C19ORF47, which decreased the expression of > 400 genes, many encoding extracellular matrix proteins with roles in cell adhesion and signalling. Genes with decreased expression upon PPIL4 knockdown were modestly enriched for roles in amino acid metabolism (**Extended Data Fig. 9d, e**).

Next, we assessed whether depleting each protein affected alternative splicing by monitoring changes in cassette exons, microexons, alternative 5′ and 3′ splice sites, and intron retention (**Fig. 5c**). PPIL4 depletion had the most widespread effect on splicing, reducing the inclusion of 864 cassette exons and increasing the inclusion of 1,246 introns, suggesting potential roles in controlling both alternative exon usage and intron retention. PAXBP1 knockdown resulted in substantial changes in alternative 5′ splice-site usage, while C19ORF47 knockdown resulted in relatively fewer changes in splicing.

Previously, through proximity labelling using APEX2-tagged nuclear speckle markers, we provided evidence that nuclear speckles are associated with functionally coordinated programs of highly-regulated introns that share hallmark features of nuclear ‘detained introns’^29^. Accordingly, we next assessed the overlap between PPIL4-regulated introns and the APEX2-nuclear speckle marker-labelled introns, and as control comparisons, a distinct set of retained introns labelled by APEX2- tagged LMNA and a group of randomly selected introns. The PPIL4-regulated introns had significantly greater overlap with the speckle-associated introns than the lamin-associated or random introns (84/1,661, *p <* 3.49 × 10^-14^, Fisher’s Exact test; **Extended Data Fig. 9f**). Consistently, they also share hallmark sequence features of speckle-associated retained introns, including relatively short length (p < 2.13 × 10^-187^, Wilcoxon test; **Fig. 5d**), having significantly higher G/C content than the random or lamina-enriched introns (p < 9.08 × 10^-291^, Wilcoxon test; **Extended Data Fig. 9g**), and having weaker 5′ and 3′ splice sites (*p* < 4.76 × 10^-24^ and *p <* 5.75 × 10^-19^, Wilcoxon test; **Extended Data Fig. 9h**). As many intron-containing transcripts are predicted to result in RNA decay^52,53^, we tested this for PPIL4- regulated transcripts by comparing their relative levels with and without introns. Although only 21/571 genes with PPIL4-regulated retained introns showed significant expression changes, minor expression decreases after PPIL4 knockdown were significantly more common in genes with PPIL4-regulated introns compared to those without (*p* = 2.513 × 10^-12^, signed-rank test; **Fig. 5e**). Together, these results reveal novel nuclear speckle-associated proteins that regulate mRNA transcript levels and alternative splicing.

### QKI is a novel core-localised paraspeckle protein that dictates paraspeckle size

We identified and validated the RNA binding protein QKI, a splicing regulator with important roles in cellular differentiation, development, and pathology^54,55^, as a paraspeckle protein. Supporting its assignment, public UV crosslinking and immunoprecipitation (CLIP) data^56^ for QKI in HepG2 cells revealed distinct binding to NEAT1_2, (NEAT1’s long isoform, the scaffolding lncRNA for paraspeckles; **Extended Data Fig. 10a**). We found that QKI colocalised with NEAT1_2 and NONO in HeLa and HEK293 cells (**Extended Data Figs. 8b** and **10b**), consistent with a previous report^57^. BioID profiling of QKI recovered many preys also recovered by essential paraspeckle proteins (**Fig. 6a**). Paraspeckles have a core-shell structure, with the 5′ and 3′ ends of NEAT1_2 located in the shell region and its central sequence in the core^58^. Different paraspeckle proteins reside in specific sub- regions. Consistent with its central NEAT1 binding site in the CLIP data, super-resolution 3D- structured illumination microscopy (3D-SIM) in HeLa cells revealed QKI in the paraspeckle core, surrounded by a NEAT1 5′-positive shell (**Fig. 6b**).

**Fig. 6.**
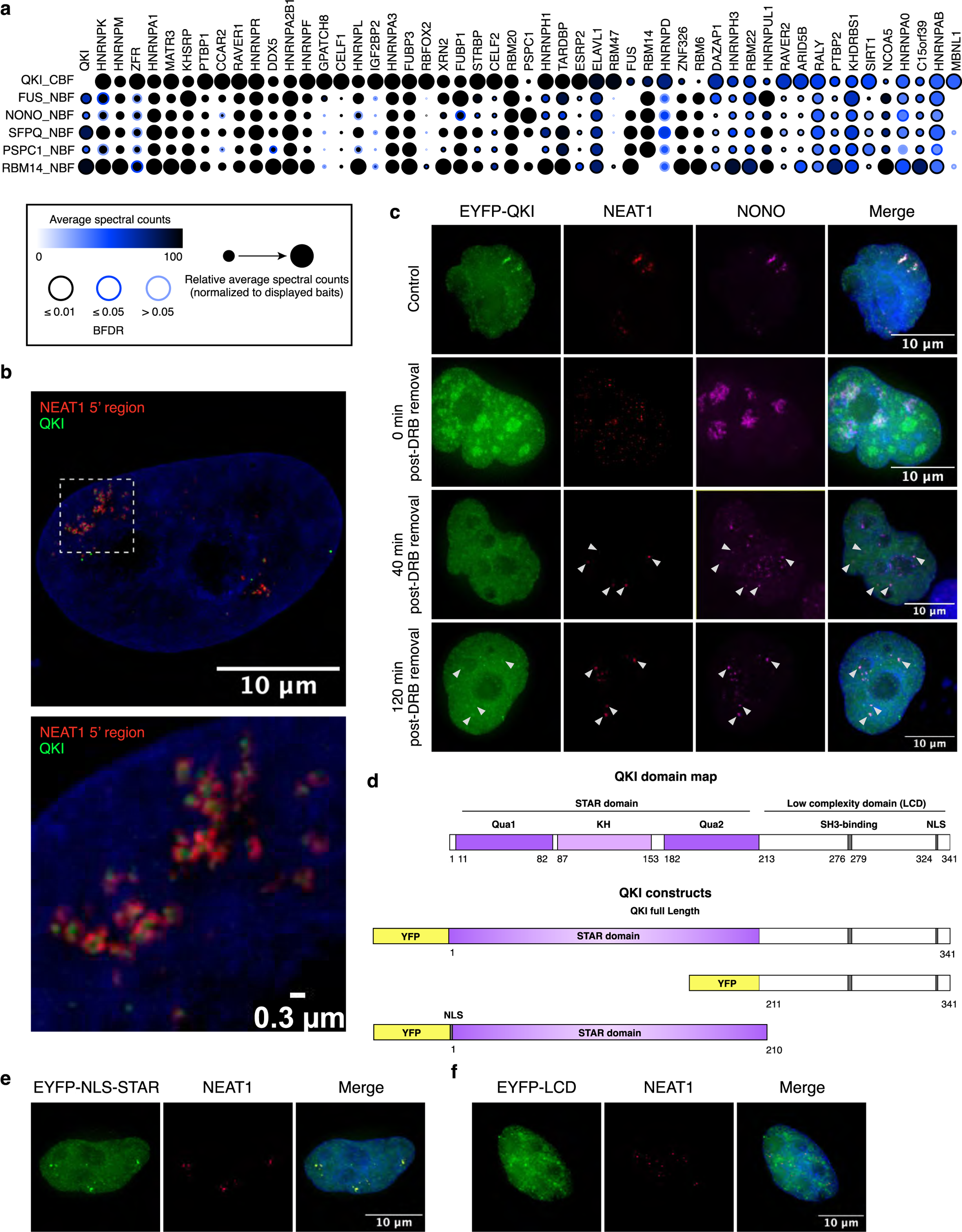
QKI is a novel core-localised paraspeckle protein. **a**, BioID data for QKI and canonical paraspeckle markers, including the core-localised proteins NONO, SFPQ, and PSPC1. CBF and NBF indicate C- and N-terminal fusions, respectively. **b**, Super-resolution 3D-SIM microscopy of QKI (green) and the 5′ region of NEAT1 (red) in HeLa cells. DAPI was used to visualize the nucleus. **c**, EYFP-QKI-expressing HeLa cells were transiently treated with DRB to induce paraspeckle disassembly. DMSO was used as a control treatment. The merged images include DAPI staining to visualize the nucleus. White arrowheads indicate paraspeckles. **d**, Schematics of QKI’s domain structure (top) and the constructs used in this study (bottom). **e**, Localisation of EYFP-NLS-STAR in HeLa cells subjected to RNA-FISH for NEAT1. DAPI was used to visualize the nucleus. **f**, Localisation of EYFP-NLS-LCD in HeLa cells subjected to RNA-FISH for NEAT1. DAPI was used to visualize the nucleus.

Given its interactions and localisation, we hypothesised that QKI regulates paraspeckle formation. Several known paraspeckle proteins (SFPQ, NONO, RBM14) are essential for NEAT1_2 stability. Others (FUS, DAZAP1, HNRNPH3, and the SWI/SNF complex) are essential for paraspeckle assembly. To examine QKI’s recruitment to nascent paraspeckles, we treated HeLa cells expressing enhanced yellow fluorescent protein (EYFP)-tagged QKI with 5,6-dichloro-1-beta-D- ribofuranosylbenzimidazole (DRB), which induces paraspeckle disassembly^35,59^, then removed the DRB and visualised QKI, NONO, and NEAT1_2 during paraspeckle reassembly. NONO colocalised with NEAT1_2 40 min after DRB removal, and QKI joined them 120 min after NONO (**Fig. 6c**). To further explore QKI’s recruitment to assembling paraspeckles, we used the proteasome inhibitor MG132, which increases paraspeckle size^16^. Immunofluorescence imaging showed that QKI was sequestered in paraspeckles after MG132 treatment (**Extended Data Fig. 10c**).

QKI contains a STAR domain (including an N-terminal Qua1 dimerisation domain, a K- homology (KH) RNA binding domain, and a C-terminal Qua2 domain) and a low-complexity domain (LCD)—a type of IDR, which we found to be highly enriched in paraspeckle proteins (**Extended Data Fig. 3c**). To determine if these regions target QKI to paraspeckles, we expressed EYFP fusions of each domain in HeLa cells to compare with a full-length fusion (**Fig. 6d**). The STAR domain alone was sufficient for paraspeckle localisation (**Fig. 6e**), while the LCD alone was insufficient (**Fig. 6f**). QKI RNA binding activity depends on homodimerisation^60^. Introducing dimerisation-preventing point mutations in the STAR domain revealed that dimerisation/RNA binding was necessary to localise both full-length QKI (**Fig. 7a**) and the isolated STAR domain (**Fig. 7b**) to paraspeckles.

**Fig. 7.**
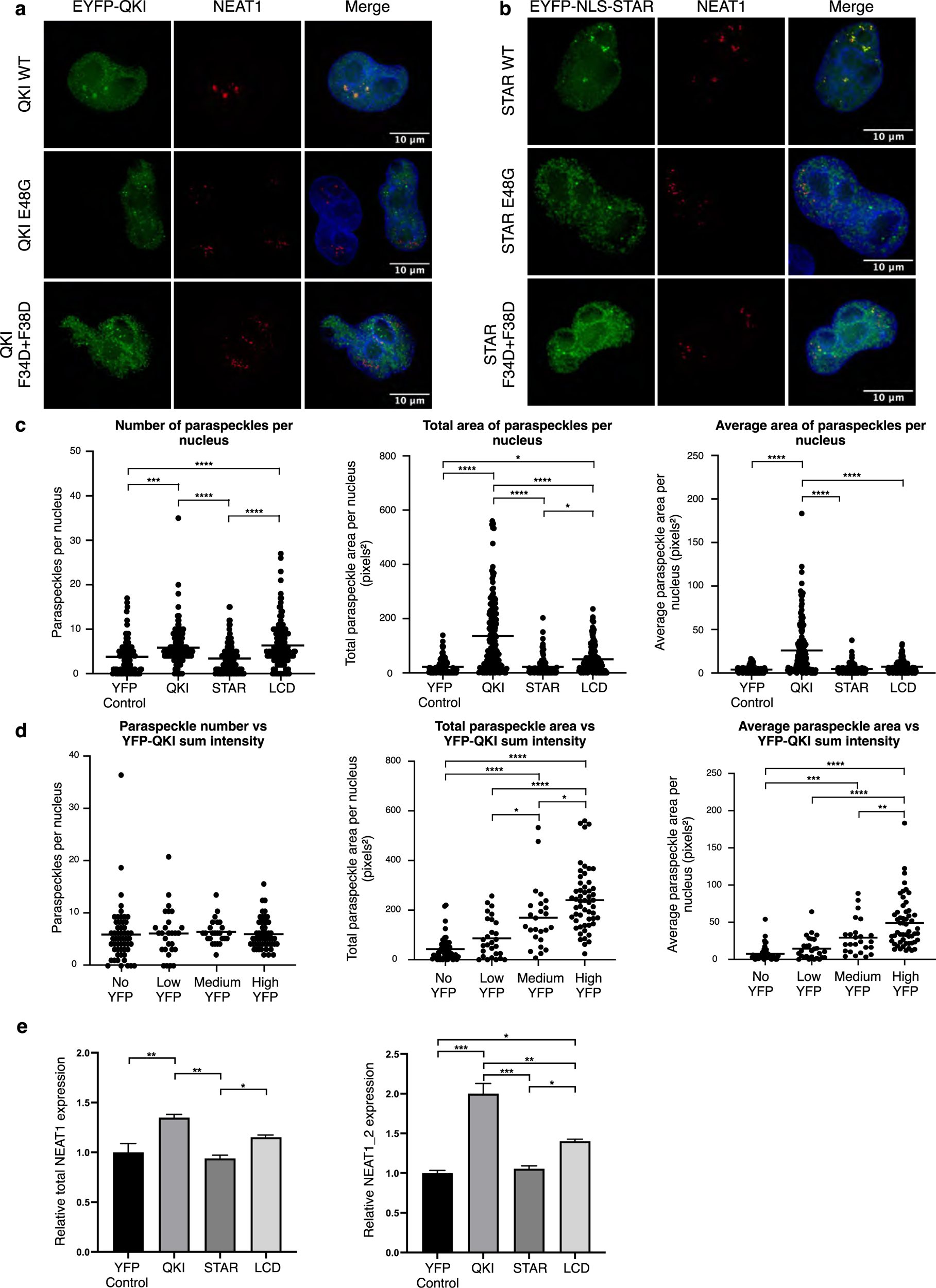
QKI dictates paraspeckle size *via* its STAR domain. **a**, Localisation of wild-type (WT) EYFP-QKI compared to STAR domain mutants that prevent homodimerisation in HeLa cells. The merged images include DAPI staining to visualize the nucleus. **b**, Localisation of WT EYFP-NLS-STAR compared to homodimerisation mutants in HeLa cells. The merged images include DAPI staining to visualize the nucleus. **c**, Quantitative microscopy of the effects of QKI (or QKI subdomain) overexpression on paraspeckle number and size (area). At least 100 nuclei were analysed for each condition. Paraspeckle numbers: ***, *p* = 0.0005, ****, *p <* 0.0001; total area: * (control vs LCD), *p* = 0.0214, * (STAR vs LCD), *p* = 0.0147, ****, *p <* 0.0001; average area: ****, *p <* 0.0001 by ordinary one-way analysis of variance (ANOVA) with Tukey’s multiple comparisons test. All other comparisons were not statistically significant. **d**, Quantitative microscopy of QKI expression (using the EYFP intensity as a proxy) and paraspeckle numbers and sizes. At least 25 nuclei per technical replicate were analysed for each condition. Average paraspeckle area vs EYFP-QKI sum intensity: ** *p* = 0.0033, *** *p* = 0.0007, **** *p <* 0.0001 by ordinary one-way ANOVA with Tukey’s multiple comparisons test. All other comparisons were not statistically significant. **e**, RT-qPCR of total NEAT1 (both isoforms) and long isoform (NEAT1_2) levels in response to QKI or QKI subdomain overexpression in HeLa cells. Total NEAT1: (control vs QKI) ** *p* = 0.0079, (QKI vs STAR) ** *p* = 0.0044, (STAR vs LCD) * *p* = 0.0451; NEAT1_2: (control vs QKI) *** *p* = 0.0005, (control vs LCD) * *p* = 0.0155, (QKI vs STAR) *** *p* = 0.0006, (QKI vs LCD) ** *p* = 0.035, (STAR vs LCD) * *p* = 0.0260 by ordinary one-way ANOVA with Tukey’s multiple comparisons test. All other comparisons were not statistically significant.

QKI overexpression significantly increased paraspeckle size, with a smaller effect on their quantity per nucleus (**Fig. 7c**). However, although the STAR domain was sufficient to target QKI to paraspeckles (**Fig. 6e**), the full-length protein was required for QKI to regulate paraspeckle size. The increase in paraspeckle size (but not their quantity) was proportional to the QKI expression level (**Fig. 7d**). QKI overexpression also significantly increased the NEAT1 level, particularly that of NEAT1_2 (**Fig. 7e**). Altogether, these data support a direct role for QKI in regulating paraspeckle size upon recruitment to NEAT1_2.

## Discussion

This comprehensive proximity labelling-based survey identified distinct clusters of proteins comprising the paraspeckle, nuclear speckle, Cajal body, PML body, and nucleolus, including over 350 novel components. We also profiled baits representing the nuclear stress body^61^, histone locus body^62^, Sam68 nuclear body^63^, polycomb body^64^, and perinucleolar compartment^65^; however, our NMF analyses did not resolve them into distinct clusters. This was expected for nuclear stress bodies, since biotin labelling was performed at steady-state (and their components are primarily annotated as nuclear speckle proteins^66^). The absence of a perinucleolar compartment cluster was also unsurprising, as these are found in cancer cells. However, known perinucleolar compartment proteins^67^ were found in our paraspeckle cluster, and PTBP1, PTBP3, and RAVER1 were validated as paraspeckle proteins. Paraspeckle proteins relocate to perinucleolar cap structures in response to transcriptional inhibition^35,59,18^; however, their functional significance remains elusive. Profiling our nuclear speckle and paraspeckle baits under dynamic, stress-induced conditions could inform on how they regulate gene expression during stress. Sam68 nuclear body baits were also strongly correlated and largely clustered with paraspeckle baits, and STRBP was validated as a novel paraspeckle protein. This suggests that, unlike in HeLa cells^63^, Sam68 nuclear bodies may be indistinct from paraspeckles in HEK293 cells. Together, our data demonstrate the highly interconnected nature of nuclear bodies.

We validated 29 novel paraspeckle and nuclear speckle components, and provided compelling preliminary evidence that three nuclear speckle preys regulate mRNA transcript levels and splicing. PPIL4 and PAXBP1 depletion caused profound and diverse splicing changes, suggesting general regulatory roles. Consistently, they were assigned to both the nuclear speckle and splicing clusters. PAXBP1 formed proximal associations with known splicing regulators, including DDX41, which was recently linked to RNA polymerase II pausing at 5′ splice sites^68^. Accordingly, PAXBP1 knockdown disproportionately affected alternative 5′ splice site usage. Conversely, PPIL4 did not associate with known splicing regulators, despite its RNA recognition motif. Instead, it recovered proteins associated with the polymerase II C-terminal domain (CTD) and involved in transcriptional termination and mRNA 3′ end processing (*e.g*., PCF11^69^ and RPRD1B^70^). In *Arabidopsis*, PPIL4 (AtCyp59) localises to puncta near active transcription sites and associates with the polymerase II CTD and RNA^71,72^. One possible hypothesis is that PPIL4 isomerizes proline-directed phosphorylation sites in the CTD and other proximal interactors to alter transcriptional kinetics or termination, which both influence splicing^73^. A similar mechanism was proposed for the peptidyl-prolyl cis-trans isomerase PIN1, which influences CTD phosphorylation^74–76^. We also identified a role for C19ORF47 in transcriptional regulation. C19ORF47 formed proximal interactions with many critical members of the transcription and export complex^77^, including ALYREF, CHTOP, FYTTD1, NCBP3, POLDIP3, and ZC3H11A, suggesting a role in promoting mRNA export. While further mechanistic experiments on PAXBP1, PPIL4, and C19ORF47 are needed to uncover their roles in transcriptional and post-transcriptional regulation, these examples highlight the power of our approach to identifying new and functionally relevant nuclear body components.

We also identified QKI, an RNA-binding protein that regulates splicing, as a novel core- localised paraspeckle protein and the first specific regulator of paraspeckle size. QKI is involved in central nervous system and cardiovascular development^54,55^ and—like paraspeckles^17^—is implicated in neurodegenerative disease^78^. Interestingly, while QKI localised to the paraspeckle core and its preys highly overlapped with those of known core proteins, its recruitment occurred later than the paraspeckle assembly-initiating protein NONO, and its overexpression affected the size of paraspeckles but not their quantity, indicating that QKI affects paraspeckle maturation rather than initial formation. We also found that QKI’s recruitment to NEAT1 requires dimerisation *via* its STAR domain. Once bound, QKI’s low complexity domain—an IDR—helps modulate paraspeckle size. IDRs, which drive phase separation^79,80^, were enriched among nuclear body components in our dataset, particularly in paraspeckles. IDRs in FUS and RBM14 are crucial for paraspeckle phase separation^81^, and our results suggest that QKI’s IDR promotes this as well. Whether other proteins are involved, including various other QKI-associated RNA-binding proteins, remains unknown. We expect that multivalent protein-protein and protein-RNA interactions are responsible and that NEAT1_2 stabilisation or upregulation is involved, as QKI overexpression preferentially increased NEAT1_2 over NEAT1_1. Our extensive paraspeckle protein BioID data and proteome (*i.e*., cluster 0) provide a wealth of starting points for continued mechanistic studies into their dynamics.

This study has a few limitations. For instance, while the paraspeckle and nucleolus are noted for having substructural organisation^58,82^, these were not resolvable here, possibly because the long (8 h) biotin labelling treatment prevented specific subcompartment labelling. Coupling proximity labelling to chemical cross-linking^83^ or using an enzyme with a faster labelling time could also help provide adequate resolution. Furthermore, including additional baits for several nuclear bodies (*e.g*., Sam68 nuclear bodies, histone locus bodies) could increase their coverage and resolve specific clusters.

As dynamic membraneless organelles, nuclear bodies have largely evaded interrogation by traditional proteomic approaches, and targeted colocalisation screens of biomolecular condensates are susceptible to morphology and localisation changes due to formaldehyde fixation^84^. Coupling BioID to mass spectrometry and NMF clustering enabled the proteomic investigation of diverse nuclear bodies, generating a rich dataset that can be explored as individual high-confidence bait-prey relationships or as clusters representing different nuclear structures. It contains a wealth of information that will accelerate our understanding of nuclear structures and processes, and provides an adaptable strategy for high-resolution profiling of the myriad subcellular domains found in mammalian cells.

## Extended data figure legends

**Extended Data Fig. 1 | Baits representing different nuclear bodies show expected staining patterns in cells and are highly correlated (related to Fig. 1)**

**a,** Immunofluorescence imaging of the expression (FLAG) and biotinylation (Streptavidin) of selected baits. The merged images include DAPI staining to visualize the nucleus. CBF and NBF indicate C- and N-terminal fusions, respectively. Additional bait images are available online at https://prohits-web.lunenfeld.ca/.

**b,** Scatter plot of the Pearson correlations between the spectral counts for each unfiltered prey in two independent biological replicates.

**c,** Heat map of the pairwise Pearson correlations between the unfiltered, pre-SAINTexpress spectral counts of the baits.

**Extended Data Fig. 2 | GSEA of baits representing each nuclear body and summary of their preys (related to Fig. 1)**

**a,** Overview of unique high-confidence (SAINTexpress BFDR ≤ 0.01) preys recovered across all baits. The colours indicate bait categories.

**b,** GSEA results for selected baits. Columns indicating enrichment in custom literature-curated terms for specific nuclear bodies are indicated in bold. Full results are provided in Supplementary Table 4.

**Extended Data Fig. 3 | Baits chosen to profile nuclear bodies recover highly overlapping sets of preys (part 1/2; related to Fig. 1)**

**a,** A network of bait-prey relationships among the top 25 high-confidence preys per bait by length- normalised spectral counts. The colours indicate bait categories.

**b,** A bait-prey subnetwork containing 7/21 paraspeckle baits and the RNA binding protein PTBP1. Large and small nodes denote baits and preys, respectively. Arrowheads indicate bait-prey relationships. Known paraspeckle proteins are shown in pink.

**c,** A bait-prey subnetwork containing 6/16 PML body baits, plus EMD and LMNB1. Large and small nodes denote baits and preys, respectively. Arrowheads indicate bait-prey relationships. Known PML body proteins are shown in teal.

**d,** A bait-prey subnetwork containing 5/15 nuclear speckle baits, plus SNPRPD3 (which was primarily chosen to profile Cajal bodies but is also associated with nuclear speckles). Large and small nodes denote baits and preys, respectively. Arrowheads indicate bait-prey relationships. Known nuclear speckle proteins are shown in blue.

**e,** A bait-prey subnetwork containing 3/5 histone locus body baits. SLBP (tagged at both termini) was not included, as it was most closely related to nucleolar proteins. Large and small nodes denote baits and preys, respectively. Arrowheads indicate bait-prey relationships. Known histone locus body proteins are shown in maroon.

**Extended Data Fig. 4 | Baits chosen to profile nuclear bodies recover highly overlapping sets of preys (part 2/2; related to Fig. 1)**

**a,** A bait-prey subnetwork containing 6/23 nucleolus baits, plus coilin (COIL), which was primarily chosen to profile Cajal bodies, but also associates with the nucleolus. Large and small nodes denote baits and preys, respectively. Arrowheads indicate bait-prey relationships. Known nucleolar proteins are shown in orange.

**b,** A bait-prey subnetwork containing 9/23 Cajal body baits plus RIOK1, which is primarily annotated as a nucleolar protein. Large and small nodes denote baits and preys, respectively. Arrowheads indicate bait-prey relationships. Known Cajal body proteins are shown in green.

**Extended Data Fig. 5 | A 19-cluster NMF analysis best recapitulates known nuclear body compositions and interconnections (related to Fig. 2)**

**a,** F-scores for the nuclear bodies of interest in analyses resulting in *k* = 10–24 clusters. The average F- scores across the highest-scoring Cajal body, nuclear speckle, nucleolus, paraspeckle, and PML body clusters are shown on the left.

**b,** F-scores for the nuclear bodies of interest in analyses with *k* = 19 clusters and primary threshold values of 0–50% of the maximum NMF score in each cluster. The average F-scores across the highest- scoring Cajal body, nuclear speckle, nucleolus, paraspeckle, and PML body clusters are shown on the left.

**c,** F-scores for the nuclear bodies of interest in analyses with *k* = 19 clusters and different multiple cluster assignment thresholds. The average F-scores across the highest-scoring Cajal body, nuclear speckle, nucleolus, paraspeckle, and PML body clusters are shown on the left.

**d,** Heat map of the Pearson correlations between the NMF scores of preys in different clusters.

**e,** Heat map of the number of preys in each cluster with multiple rank assignments.

**Extended Data Fig. 6 | Top 50 preys by NMF score in the Cajal body, nucleolus, and PML body clusters (related to Fig. 3)**

**a,** A hierarchical clustering of the NMF scores of the top 50 genes in the Cajal body cluster (10). Known Cajal body proteins are in green, and baits in this study are indicated in bold.

**b,** A hierarchical clustering of the NMF scores of the top 50 genes in the nucleolar cluster (4). Known nucleolar proteins are in green, and baits in this study are indicated in bold.

**c,** A hierarchical clustering of the NMF scores of the top 50 genes in the PML body cluster (14). Known PML body proteins are in green, and baits in this study are indicated in bold.

**Extended Data Fig. 7 | Nuclear body-associated NMF clusters are enriched for specific domains and the propensity to phase separate (related to Fig. 3)**

**a,** Dot plot of the top enriched protein domains in the NMF clusters. Domains and clusters are shown in alphabetical and numerical order, respectively. Full results are provided in Supplementary Table 5.

**b,** Violin plot of the distributions in predicted phase separation propensity for the proteins in each cluster, arranged by descending median score. The entire human proteome, the HEK293 proteome, all high-confidence preys (BFDR ≤ 0.01) in this dataset, and all high-confidence preys from two other BioID datasets (the human cell map (BFDR ≤ 0.01) and stress granule/P body map (AvgP ≥ 0.95) are also shown. The red dashed line indicates the threshold for proteins with the highest confidence predicted phase separation propensity.

**Extended Data Fig. 8 | Novel nuclear speckle and paraspeckle proteins recover known components of their respective nuclear bodies (related to Fig. 4)**

**a,** Representative fluorescence images of validated paraspeckle proteins in HeLa cells. The merged images include DAPI staining to visualize the nucleus.

**b,** Representative fluorescence images of validated paraspeckle proteins in HEK293 cells. The merged images include DAPI staining to visualize the nucleus. White arrowheads indicate colocalised foci.

**c,** Bar plot of the total preys in the paraspeckle and nuclear speckle NMF clusters recovered by each reciprocal BioID bait (related to the data in Fig. 4e). CmT and NmT indicate C- and N-terminal miniTurbo fusions, respectively.

**Extended Data Fig. 9 | Nuclear speckle proteins regulate gene expression and alternative splicing (related to Fig. 5)**

**a,** Domain maps of the three proteins profiled (RRM, RNA-recognition motif).

**b-c**, Relative changes in PPIL4, PAXBP1, and C19ORF47 protein (b) and transcript (c) abundance following siRNA knockdown.

**d-e,** Enriched GO terms among genes with decreased expression following C19ORF47 (d) or PPIL4 (e) knockdown (from Fig. 5b).

**f,** Percentage of PPIL4-regulated retained introns which are speckle-associated, lamina-associated (Barutcu, Wu *et al*. 2022, PMID: 35182477), or random introns. 84/1,661 introns with increased retention in response to PPIL4 knockdown are speckle-associated (*p <* 3.49 × 10^-14^, Fisher’s Exact test).

**g-h**, GC content (g) and predicted 5’ and 3’ splice site strength scores (h) for PPIL4-retained introns compared to speckle-/lamina-associated introns from (Barutcu, Wu *et al*. 2022, PMID: 35182477) or a randomly selected group.

**Extended Data Fig. 10 | QKI can interact with NEAT1 and colocalises with paraspeckle markers (related to Fig. 6)**

**a,** Public CLIP data (ENCODE identifier ENCSR570WLM) for NEAT1_2. The QKI binding motif is indicated.

**b,** Fluorescence microscopy of endogenous QKI and paraspeckle markers in fixed HeLa cells. The merged image includes DAPI to visualize the nucleus.

**c,** Fluorescence microscopy images of pEYFP-QKI transfected HeLa cells treated with 5 µM MG132 for 17 h to induce paraspeckle growth. The merged images include DAPI to visualize the nucleus.

## Supplementary information

**Figures**

**Supplementary Fig. 1 | Additional fluorescence microscopy results for paraspeckle and nuclear speckle candidate validation**

**Tables**

**Supplementary Table 1 | Literature-curated lists of known nuclear body proteins**

This table includes the referenced curated lists of nuclear body proteins, tables sourced from previous publications, and custom g:Profiler tokens and .GMT file source data for these terms.

**Supplementary Table 2 | Baits used for BioID (BirA* map)**

This table lists the initial 146 baits selected, their tag positions, the compartment or function they were chosen to profile, and additional relevant information.

**Supplementary Table 3 | SAINT results for the BirA* dataset**

Full bait-prey data and SAINTexpress for the main dataset. This corresponds to 7013_cleaned_v2.txt found in the additional materials associated with this manuscript

**Supplementary Table 4 | Gene set enrichment analysis results for preys recovered by the baits chosen for each nuclear body**

Related to Extended Data Fig. 2b. GSEA of the preys recovered by each bait, organized by bait category. Tabs for GSEA using g:Profiler including custom literature-curated terms, indicated with “CUSTOM”. Other tabs contain only cellular compartment, molecular function, and biological function GO terms.

**Supplementary Table 5 | NMF results**

This table contains the bait matrix, prey matrix, full gene lists for the clusters, GSEA results from g:Profiler, statistically significant differences in IDR composition between clusters, and PFAM domain enrichment analysis results. Related to Fig. 2–3 and Extended Data Fig. 5–7.

**Supplementary Table 6 | Baits used for reciprocal BioID**

This table lists all 40 validation baits with their tag, tag position, isoform/source, and rationale for selection. This includes the 20 validation candidates described in Fig. 4 and Extended Data Fig. 8, including their predicted structures and validation results.

**Supplementary Table 7 | SAINT results for the miniTurbo dataset**

Full bait-prey data and SAINTexpress scoring for the reciprocal BioID experiments. Related to Fig. 4c and Extended Data Fig. 8c. This corresponds to 7015_final.txt found in the additional materials associated with this manuscript.

**Supplementary Table 8 | Comprehensive validation results**

This table lists the NMF scores of candidates tested by fluorescence microscopy colocalisation or reciprocal BioID, plus associated validation results.

## Methods

### Data availability

The BioID data have been deposited in the MassIVE repository (https://massive.ucsd.edu/ProteoSAFe/static/massive.jsp). The BirA* dataset was assigned the accession number MSV000090684. The ProteomeXchange accession number is PXD038077. The miniTurbo dataset was assigned the accession number MSV000094294. The ProteomeXchange accession number is PXD050558. All RNA-Seq data and analyses have been deposited in the Gene Omnibus (GEO: GSE264355).

A searchable version of the BirA* dataset is available at https://prohits-web.lunenfeld.ca/ (data set 39). In addition to enabling exploration of all high-confidence (or all detected) proximity interactions, we also display Cytoscape network images that organize the top recovered preys for each bait for convenient examination. Immunofluorescence imaging data for baits (as shown in Extended Data Fig. 1a) is also hosted at this site.

### Code availability

The code used for analysis and data visualisation in this manuscript are available on GitHub (https://github.com/gingraslab/Dyakov-et-al-2024).

### Contact for Reagent and Resource Sharing

Further information and requests for resources and reagents should be directed to the Lead Contact, Anne-Claude Gingras.

## Method details

### BioID bait selection

We selected baits for BioID by comparing our curated list of known nuclear body proteins (**Supplementary Table 1**) with locally available ORFs. We also profiled 36 baits (representing 33 unique genes) previously profiled in Go *et al.* 2021^24^, Youn *et al.* 2018^28^, or other collaborative studies^85,86^ that reliably recover components of nuclear structures or pathways related to RNA metabolism. In total, we profiled 146 baits representing 123 unique genes (21 baits were tagged at both termini, and three C-terminally tagged isoforms were profiled for HNRNPR; **Supplementary Table 2**).

### Stable cell line generation, bait induction, labelling, and harvest

Flp-In T-REx HEK293 cells were transfected using jetPRIME transfection reagent using the manufacturer’s protocol with minor modifications: cells were seeded in 6-well plates and cultured in 2 mL Dulbecco’s modified Eagle’s medium (DMEM) supplemented with 5% fetal bovine serum (FBS), 5% Cosmic Calf Serum, and 100 U/mL Pen/Strep. Cells were transfected at 70–80% confluency with ∼100 ng of a pcDNA5 plasmid expressing a BirA*-FLAG- or miniTurbo-FLAG-tagged bait and 1 µg of pOG44 with 2 µL of jetPRIME reagent in 200 µL of jetPRIME buffer. The cells in each well were transferred to a 10 cm dish the next day, and selected with hygromycin (200 µg/mL final concentration) starting the day after that. The selection medium was replaced every 2–3 days until visible hygromycin- resistant colonies formed. The colonies for each bait were then pooled in a 15 cm dish and cultured to 75–85% confluency. BirA*-FLAG-tagged baits were induced for 17 h by replacing the medium with fresh medium (without hygromycin) and 1 µg/mL tetracycline, then 50 µM biotin was added for 8 h.

Baits fused to miniTurbo were induced for 23 h and biotin labelled for 1 h. After biotin labelling, the DMEM was aspirated and the cells were rinsed with ice-cold PBS, harvested in 1 mL of ice-cold PBS with a cell scraper, and transferred to a pre-weighed 2 mL tube. The cells were pelleted by centrifugation at 500 × *g* for 5 min at 4°C, the supernatant was aspirated, and the tubes were placed in dry ice. Following flash-freezing, the samples were weighed and stored at -80°C. Pellets typically weighed 0.1–0.2 g.

### Streptavidin affinity purification and peptide preparation

Frozen pellets were thawed on ice and lysed in modified RIPA buffer (50 mM Tris (pH 7.5), 150 mM NaCl, 1.5 mM MgCl_2_, 1 mM ethylenebis(oxyethylenenitrilo)tetraacetic acid (EGTA), 0.1% sodium dodecyl sulfate (SDS), and 1% IGEPAL CA-630), supplemented with 0.4% sodium deoxycholate and protease inhibitors (Sigma-Aldrich P8340) just before lysis. We used 400 µL buffer for each 0.1 g of cells and incubated them for 20 min at 4°C with gentle rotation (all subsequent 4°C incubations also involved gentle rotation). Samples were sonicated on a Qsonica sonicator with a CL-18 probe at 25% amplitude for three cycles (5 s on, 3 s off), then incubated with 375 U of Benzonase for 15 min at 4°C. Additional SDS was added to a final concentration of 0.4% and the lysates were incubated for 15 min at 4°C. Samples were centrifuged at 16,000 × *g* for 20 min at 4°C, and the cleared lysate (supernatant) was transferred to a new tube. Streptavidin Sepharose beads (GE 17-5113-01) were washed three times with modified RIPA buffer with 0.4% SDS and added to each tube (in 30 and 20 µL bed volumes for BirA* and miniTurbo samples, respectively) for a 3-h incubation at 4°C. Samples were washed once with wash buffer (2% SDS, 50 mM Tris (pH 7.5)), twice with modified RIPA buffer with 0.4% SDS, and three times (BirA* samples) or four times (miniTurbo samples) with 50 mM ammonium bicarbonate (pH 8.5). The tubes were centrifuged at 500 × *g* for 30 s, the supernatants were removed, and the purified proteins were digested to peptides on-bead with 1 µg trypsin in 50 mM ammonium bicarbonate (pH 8.5) overnight at 37°C with rotation. The next day, each sample was supplemented with 0.5 µg trypsin and incubated for 2 h at 37°C. Samples were gently vortexed and centrifuged at 500 × *g* for 30 s and the supernatants were transferred to new tubes. The beads were washed twice with 30 µL of high-performance liquid chromatography (HPLC)-grade water and the supernatants were added to the new tubes. Fresh 50% formic acid was added to a final concentration of 2%, then the samples were dried by vacuum centrifugation and stored at -80°C.

### Sample acquisition using TripleTOF mass spectrometers

Aliquots of each sample (6 μL in 2% formic acid; corresponding to 1/6th of the cellular material obtained from one 15-cm tissue culture plate) were directly loaded onto an equilibrated HPLC column (made in-house) at 800 nL/min. Peptides were eluted over a gradient generated by an Eksigent ekspert nanoLC 425 (Eksigent, Dublin, CA, USA) nano-pump and analysed on a TripleTOF 6600 System (AB SCIEX, Concord, ON, Canada). The gradient (2–35% acetonitrile in 0.1% formic acid) was delivered at 400 nL/min over 90 min, followed by 15 min of 80% acetonitrile in 0.1% formic acid, then a 15-min equilibration period back to 2% acetonitrile with 0.1% formic acid, for a total runtime of 120 min. To minimise carryover between subsequent samples, the analytical column was washed for 2 h between each sample, using an alternating sawtooth gradient of 35% and 80% acetonitrile (both in 0.1% formic acid) at 1,500 nL/min, holding each concentration for 5 min. Analytical column and instrument performance were verified after each sample by loading 30 fmol bovine serum albumin (BSA) tryptic peptide standard with 60 fmol α-casein tryptic digest and running a short 30 min gradient. TOF-MS mass calibration was performed on BSA reference ions before running the next sample to adjust for mass drift and verify peak intensity. Each sample was independently injected twice, enabling analysis in both data-dependent acquisition (DDA) and data-independent acquisition (SWATH) modes. The DDA method consisted of one 250-ms MS1 TOF survey scan (400–1800 Da) followed by ten 100-ms MS2 candidate ion scans (100–1800 Da) in high sensitivity mode. Only ions with charges of 2+ to 5+ that exceeded a threshold of 300 counts per second were selected for MS2, and former precursors were excluded for 7 s. The SWATH method consisted of one 250-ms MS1 scan followed by 54 isolation windows between 400 and 1250 amu, with a 3.8-s cycle time) The SWATH windows had variable widths (7–50 Da) and overlapped by 0.5 Da.

### SWATH-MS analysis

SWATH MS data were analysed using MSPLIT-DIA^87^ (version 1.0) implemented in ProHits 4.0^88^. To generate a sample-specific spectral library, peptide-spectrum matches from matched DDA runs were pooled by retaining only the spectrum with the best MS-GFDB (Beta version 1.0072 (6/30/2014)^89^) probability for each unique peptide sequence and precursor charge state, and a peptide-level false discovery rate (FDR) of 1% was enforced using a target decoy approach^90^. The MS-GFDB parameters were set to search for tryptic cleavages, with a precursor mass tolerance of 50 ppm and charges of 2+ to 4+. Peptide length was limited to 8–30 amino acids, and oxidized methionine was selected as a variable modification. The spectra were searched against the NCBI RefSeq database (version 57, January 30th, 2013), containing 36,241 human and adenovirus sequences supplemented with common contaminants from the Max Planck Institute (http://141.61.102.106:8080/share.cgi?ssid=0f2gfuB) and the Global Proteome Machine (http://www.thegpm.org/crap/index.html). The spectral library was then used to match peptide spectra to proteins via MSPLIT as previously described^89^. MSPLIT-DIA-identified peptides passing the 1% FDR threshold were subsequently matched to genes in ProHits. MSPLIT searches used a parent mass tolerance of ± 25 Da and a fragment mass tolerance of ± 50 ppm. When a retention time was available within the spectral library, a threshold of ± 5 min was applied to spectral matching, as done previously^89^.

### SAINT analysis

SAINTexpress (version 3.6.1)^33^ was used to score proximity interactions in the MSPLIT-DIA data. For each prey identified by a given bait, SAINTexpress calculates the probability of a true proximity interaction by comparing the spectral counts of the prey (as a proxy for abundance) relative to in negative control runs. The MSPLIT results were filtered to remove any preys identified with fewer than 2 unique peptides, and shared peptide hits were merged. Each bait was analysed in two independent biological replicates and compared to 40 negative control runs (20 BirA*-FLAG-only samples (controlling for the biotinylation caused by an untethered BirA* expressed at relatively high levels compared to the baits) and 20 untransfected parental cell line samples (controlling for both endogenous biotinylation and non-specific binding to the Sepharose). These controls were compressed to 10 virtual controls to maximize the scoring stringency. Preys with passing a 1% BFDR filter (a Bayesian estimation based on the distribution of the average SAINT score across both biological replicates) were considered high-confidence proximity interactions (elsewhere described as a BFDR ≤0.01 threshold). Non-human Uniprot IDs were manually removed, and outdated gene symbols for baits and preys were manually updated to current HGNC symbols. Six baits were flagged as poor-quality samples (due to misfolding and/or mislocalisation) and removed from the dataset after SAINT analysis (NTRK3, IQCN, SMARCA4, and C-terminal fusions of SYNE3, SRSF2, and SUMO1). Specific bait results were visualized as dot plots (*e.g*., Fig. 6a) using ProHits-viz^91^ (prohits-viz.org). In these dot plots, once a prey passes the 1% FDR threshold with at least one bait, its quantitative values are indicated for all baits shown, with their significance indicated by the dot’s border colour. In Fig. 6a, high-confidence preys recovered by QKI were sorted in descending order, and the top 51 (of 81) were plotted. The SAINT results in Supplementary Tables 3 and 7 are also provided as .txt files in the extended materials for this manuscript; these files can be uploaded to ProHits-viz for additional analysis/visualisation of bait-prey data as dot or prey specificity plots.

### Reproducibility metrics

To assess the reproducibility of our independent biological replicates for BioID analysis, we extracted both the high-confidence (BFDR ≤ 0.01) and unfiltered prey spectral counts from our SAINT results file and calculated Pearson correlation coefficients (*R*) and coefficients of determination (*R*^2^) using a custom Python script.

### Control-subtracted length-adjusted spectral counts (NormSpec)

The prey proximity order for each bait was determined from the prey’s control-subtracted length- adjusted spectral counts (NormSpec). This was calculated by first subtracting the average spectral counts of prey *j* in control samples (using the top 10 highest counts across all controls) from its average spectral counts with bait *i*, then multiplying by the median length of all bait *i*’s preys (retrieved from UniProt) divided by the length of prey *j*.

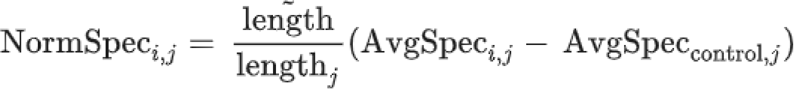

### Bait-bait correlation analyses

Bait-bait pairwise Pearson correlations were calculated based on their preys’ spectral counts in BioID experiments. No significance filter was applied, and the average of the top 10 control counts per prey was subtracted from the average spectral counts for each bait-prey association. For network visualisation using Cytoscape^92^, we filtered for correlations > 0.5, resulting in a network containing 114/140 bait nodes that highlighted the higher correlations between related baits. We used the Edge- weighted Spring Embedded layout, using the correlations as normalised value weights. The minimum, maximum, and default edge weights were set as 0.5, 1.0, and 0.68 (the average correlation among the selected associations), respectively. We used an average of 40 iterations per node and the following parameters: spring strength = 10, spring rest length = 60, disconnected spring strength = 0.5, disconnected spring rest length = 500, strength to avoid collisions = 100, layout passes = 2. For heatmap generation, all unfiltered bait-bait correlations were hierarchically clustered using complete linkage and Euclidean distance.

### Gene set enrichment analysis

GSEA was performed with g:Profiler^93^ using the default parameters and custom tokens including our curated lists of nuclear body components: gp 2aO3_xfUn_hEA (all terms as one unified source, best used when accessing g:Profiler via API call in R or Python) and gp AS1E_EjlL_7gw (terms organised into separate sources; best used when accessing g:Profiler through its web interface). For the heatmaps in Fig. 2b and 2c and Extended Data Fig. 2a, g:Profiler was run using a custom R code, then the top five highlighted driver terms were manually identified using the g:Profiler web interface. We limited our results to GO cellular compartment and biological process terms^94,95^ containing 25–500 proteins, then manually added the results for our literature-curated nuclear body terms and removed redundant terms. The results were hierarchically clustered and visualized as heat maps by ProHits-viz to determine the term order. See Supplementary Table 4 for the full enrichment results. The enrichment results for NMF cluster proteins (using a custom Python code) are provided in Supplementary Table 5.

### Bait-prey network generation

To visualise bait-prey associations as networks in Cytoscape, we first calculated control-subtracted length-adjusted spectral counts (NormSpec) for each high confidence (BFDR ≤ 0.01) bait-prey relationship. To account for baits profiled with multiple tag locations or isoforms (and to enable those bait nodes to be considered target nodes when they are preys of different baits in the dataset), we merged all parent bait genes and used the average of their prey NormSpec results when both baits recovered the same prey. The top 25 preys per bait by NormSpec were included in the Edge-weighted Spring Embedded Layout-generated network, with their NormSpec values used as normalised value edge weights. The minimum, maximum, and default edge weights were 1.17, 8161, and 269, respectively, corresponding to the minimum, maximum, and average NormSpec values of the top 25 preys. We used an average of 40 iterations per node and the following parameters: spring strength = 10, spring rest length = 60, disconnected spring strength = 0.5, disconnected spring rest length = 500, strength to avoid collisions = 100, layout passes = 2. For subnetworks, each bait node was selected along with outgoing first-neighbour bait and prey nodes and all connecting edges between the selected nodes. For visual clarity, we then applied the Prefuse Force Directed OpenCL Layout to each subnetwork, using default settings and the NormSpec values as edge weights.

### NMF analysis

We conducted NMF analysis as described in Go *et al*.^24^ with the following differences: our input matrix was the BirA* dataset with dimensions of 1,816 × 140 representing the 140 baits and 1,816 high-confidence preys (BFDR ≤ 0.01). The average of the top 10 control counts per prey was subtracted from the average spectral counts for each prey prior to analysis. The number of clusters was set as *k* = 19 based on precision-recall analyses of known proteins in each nuclear body (see our curated lists, Supplementary Table 1) for NMF runs with *k* = 10–24 (see details below). Preys were assigned to their highest scoring (*i.e.*, primary) clusters. Preys with a primary score below 5% of the maximum score within each cluster were excluded from further analysis to ensure only significant associations were considered. This means that if a prey’s strongest association with any cluster was weak, it was not included in the results. For secondary and tertiary cluster assignments, preys were included if their scores in those clusters were at least 70% of their score in their primary cluster and also met the 5% maximum cluster score threshold. Heat maps showing the top 50 preys by NMF score for selected nuclear body clusters (Fig. 3a,b and Extended Data Fig. 6a-c) were generated using ProHits-viz and hierarchically clustered (Euclidean distance, complete linkage).

### Prey-prey NMF score correlation analysis and network generation

NMF clusters were visualised in Cytoscape by first calculating pairwise prey-prey Pearson correlations based on their NMF scores. Prey pairs with correlations ≥ 0.6 were considered interactors for network- building purposes. We used this threshold because it produced robust network topology, distinctly separating each cluster while indicating their connections. The Edge-weighted Spring Embedded Layout algorithm was applied, with the correlations used as normalised value weights. The minimum, maximum, and default edge weights were set as 0.6, 1.0, and 0.85 (the average correlation between the selected prey pairs). We used an average of 40 iterations per node and the following parameters: spring strength = 15, spring rest length = 45, disconnected spring strength = 0.05, disconnected spring rest length = 2000, strength to avoid collisions = 0, layout passes = 2. Prey nodes were colour-coded by primary cluster assignment.

### Precision-recall analysis to determine the NMF cluster number and thresholds

To determine the optimal number of clusters for NMF analysis, *k* clusters from 10 through 24 were generated by NMF and compared against our curated lists of nuclear body proteins. F-scores were calculated as **2 × (precision) × (recall) / (precision + recall)** on a per-cluster, per nuclear body basis. Precision is the number of known nuclear body proteins in the cluster divided by its total number of proteins, and recall is the number of known nuclear body proteins in the cluster divided by the total number of known proteins for that nuclear body. The average F-scores across the highest scoring clusters for the Cajal body, nuclear speckle, nucleolus, paraspeckle, and PML body were used to determine the optimal number of clusters for NMF analysis, along with manual inspection to ensure clear separation of proteins known to associate with different cellular locations or processes (by GSEA of GO terms). This analysis was repeated for multiple cluster assignments and minimum NMF score thresholds with 19 clusters. We used a per-prey, per cluster threshold of 5% of the maximal score for the cluster. Preys were assigned to secondary and tertiary clusters when their score for that cluster was at least 70% of their primary (*i.e*., highest) score across all clusters.

### Domain enrichment analysis

Protein domains were retrieved from PFAM^96^ and the proteins in each cluster were tested for enrichment using a custom Python script. Domain enrichments were plotted across all clusters if they had an adjusted *p*-value < 0.01 (by Fisher’s exact test and the Benjamini-Hochberg correction) and were present in > 3 proteins in at least one cluster. The dot plot was generated in ProHits-viz^91^.

### Analysis of intrinsically disordered region composition and predicted phase separation propensity

For each protein in each cluster, we assessed the percentage of their length predicted to be intrinsically disordered and for their predicted propensity to phase separate. The 19 clusters were also compared against the entire list of 1,816 high-confidence preys, the high-confidence preys in the human cell map^24^ (4,424 proteins) and stress granule and processing body^28^ BioID datasets (4,424 and 1,792 proteins, respectively), the entire human proteome (20,300 proteins), and the HEK293 proteome (10,750 proteins). The human and HEK293 proteomes were predicted using RNA-Seq data from The Human Protein Atlas version 16.1 and Ensembl version 83.38^97^. Genes with ≥ 2.5 transcripts per million in HEK293 cells were included in the predicted proteome. Protein sequences were retrieved from UniProt (https://www.uniprot.org/proteomes/UP000005640) and IDRs were predicted using Metapredict^98^. Differences in IDR percentages between clusters were analysed using the Mann- Whitney U test (Supplementary Table 5). To analyse predicted phase separation propensity, we used LLPhyScore^40^ to predict 8-feature sum scores for each protein, in which the top 10% of scores represent high-confidence predicted phase separating proteins (see methods in Millar *et al.* 2023^99^). All analyses and plot generation were performed using custom Python codes.

### Analysis of reciprocal BioID results

Novel candidate paraspeckle- or nuclear speckle-associated proteins were fused to the miniTurbo biotin ligase for reciprocal BioID as described above. The average spectral counts of paraspeckle and nuclear speckle proteins (NMF clusters 0 and 12, respectively) were plotted for each reciprocal bait. Average spectral counts across all control samples were subtracted from the average spectral counts for each prey, and the total spectral counts for preys were normalised across baits, relative to the maximum average spectral counts for that prey across all baits. Significant differences in the per-bait distribution of recovered paraspeckle and nuclear speckle proteins were evaluated by Wilcoxon signed-rank test.

### Fluorescence colocalisation imaging

#### BioID fluorescence quality control

Flp-In T-REx HEK293 cells stably expressing BirA*-FLAG-tagged baits were grown to ∼50% confluency on 12 mm poly-L-lysine (PLL)-coated coverslips (Corning, 354085) in 12-well trays using the same conditions described for BioID-MS experiments, including overnight induction with 1 µg/mL tetracycline and 8 h of labelling with 50 µM biotin. The cells were washed with cold phosphate- buffered saline (PBS) containing 200 μM CaCl_2_ and 100 μM MgCl_2_, fixed for 10 min with 4% formaldehyde in PBS plus CaCl_2_ and MgCl_2_ at room temperature, then washed twice with PBS. The samples were then permeabilised for 10 min in 0.25% Triton X-100 in PBS, washed with PBS, and blocked in 2.5% BSA (w/v) in PBS at room temperature for 1 h. Samples were then incubated with streptavidin-coupled Alexa Fluor 488 (1:2,500, Invitrogen, S32354) in 2.5% BSA in PBS containing 4′,6-diamidino-2-phenylindole (DAPI; 1:10,000) at room temperature in the dark in a humidified chamber (these conditions were used in all following steps, unless otherwise noted). Then, samples were washed twice with 0.05% Triton X-100 in PBS and blocked in 2% skim milk powder in PBS for 1 h at room temperature. Samples were incubated with anti-FLAG M2 antibodies (1:2,000, Sigma Aldrich, F3165) in 2% milk in PBS for 1 h at 4°C, washed three times with 0.05% Triton X-100 in PBS, and incubated with Alexa Fluor 555-conjugated goat anti-mouse antibodies (1:1,000, Invitrogen, A21422) for 1 h at 4°C. After three washes with 0.05% Triton X-100 in PBS and one with PBS, coverslips were mounted on glass slides using 6 μL of ProLong Gold Antifade Mountant (Thermo Fisher Scientific, P36930). Samples were then cured overnight in the dark at room temperature, the coverslip edges were sealed with nail polish, and the slides were stored in the dark at 4°C. Images were acquired on a Nikon C1Si Confocal Microscope using a 60× objective lens and 3× field zoom, and processed using ImageJ.

#### Validation of EGFP-tagged paraspeckle and nuclear speckle candidates in HEK293 cells

Untransfected parental Flp-In T-REx HEK293 cells were grown on 12-mm PLL coverslips in 12-well trays. Validation target protein ORFs were cloned into pcDNA5-EGFP constructs and 100 ng of the expression plasmid was transiently transfected with 650 ng of carrier DNA (a pcDNA3-3×HA construct) using 1.8 µL of JetPRIME transfection reagent (Polyplus CA89129-924) in 125 µL of JetPRIME buffer. After ∼2 h, fresh medium containing 1 µg/mL tetracycline was supplied for transient overnight expression of EGFP-tagged proteins.

For RNA-FISH to examine paraspeckle colocalisation, cells grown on coverslips were fixed in cold 4% formaldehyde in PBS for 10 min, washed three times with PBS, and then permeabilised in 70% ethanol overnight at 4°C. RNA-FISH was performed against NEAT1_2 using NEAT1 Middle Segment with Quasar® 570 Dye (Stellaris FISH probes). After removing the ethanol, coverslips were incubated in 1 mL of wash buffer A (Biosearch Technologies) for 5 min. Coverslips were then placed cell side down on 50 µL of hybridisation buffer containing the probe (125 nM) in a humidified chamber and incubated in the dark at 37°C for 16 h. The coverslips were then transferred back to the 12-well tray (cell side up), washed twice in 1 mL of wash buffer A, and incubated in wash buffer A at 37°C in the dark for 30 min. Coverslips were incubated in buffer B (Biosearch Technologies) containing DAPI for 5 min at room temperature, washed once with buffer B, and then mounted onto glass slides using 6 μL of ProLong Gold Antifade Mountant. Samples were then cured, sealed, and stored as above. Z-stacks were acquired using a Delta Vision Elite wide-field microscope with a 60× objective, then subjected to 3D deconvolution and maximum intensity projection to generate single images for analysis.

Immunofluorescence to examine colocalisation was performed as described above for BioID fluorescence quality control experiments, with the following changes: an anti-SRRM2 primary antibody was used to label nuclear speckles (1:3,300, Abcam, ab11826), followed by secondary staining with an Alexa Fluor 555-conjugated goat anti-mouse antibody (1:1,000, Invitrogen, A21422). For some experiments, cells were treated with 5 µg/mL of actinomycin D for 2 h prior to fixation to increase the size of nuclear speckles. Selected samples were first subjected to RNA-FISH for NEAT1_2, followed by immunostaining for SRRM2.

## Methods related to knockdown and RNA-Seq of new nuclear speckle factors

### Knockdown and RNA isolation

For knockdown experiments, HEK293 cells were grown for 24 h in a 6-well dish, then transfected with 10 nM siRNA pools (M-018555-00-0005 (C19ORF47), M-012984-01-0005 (PAXBP1), M-010401-00-0005 (PPIL4), and D-001206-13-05 (non-targeting control); Dharmacon) using RNAiMax (Life Technologies) as recommended by the manufacturers. The medium was changed after 24 h and the cells were harvested after 72 h.

RNA was extracted using the RNeasy Mini Kit (QIAGEN) including on-column DNase digest, according to the manufacturer’s protocol. RT-qPCR reactions to measure knockdown efficiency were performed using the Maxima H Minus First Strand cDNA Synthesis Kit (Thermo) and the SensiFAST SYBR No-ROX Kit (Meridian Bioscience, BIO-98005). The 10-µL reactions contained 2 µL of the diluted cDNA template, 500 nM primers, and 5 µL of SensiFAST SYBR No-ROX, and were analysed using a CFX96 Real-Time PCR Detection System (BIO-RAD) with 2-step cycling condition with 60°C annealing temp as recommended by the manufacturer. Parallel samples were lysed in 100 mM Tris- HCl pH 7.5, 150 mM NaCl, 1% NP-40, 10% glycerol, and protease inhibitors cOmplete mini protease inhibitor cocktail (at one tablet per 10 mL lysis buffer, Roche) and western blotted for PAXBP1 (Proteintech 21357-1-AP) and C19ORF47 (Invitrogen PA5-59751) to assess knockdown at the protein level. Tubulin (Sigma #T6074) was used as a loading control. RNA-Seq was performed in biological triplicate at the Donnelly Sequencing Centre using 1 µg total RNA per sample. Sequencing libraries were prepared using Illumina’s TruSeq stranded mRNA sequencing kit using the manufacturer’s protocol modified to yield average-size fragments at ∼540 nt and sequenced using the NovaSeq S1 platform using 2×150 cycles of paired-end sequencing to a target depth of 50 million reads per sample.

### Differential expression and splicing analyses

RNA-Seq data were assessed for changes in both gene expression and splicing. For gene expression analyses, reads were quantified by pseudo-aligning pre-trimmed reads to GENCODE v43 transcripts using Salmon v1.10.1^100^ and aggregated per gene using the R package tximport^101^. Differential expression was evaluated using the classical approach (exactTest) in edgeR^102^. For each comparison, genes with a log_2_ fold change ≥ 1.0 (2-fold) and an FDR ≤ 0.05 was considered differentially expressed. Genes expressed a low levels (< 5 reads per kilobase per million mapped reads) in both the knockdown and non-targeting control samples were removed from the analysis.

For splicing analyses, RNA-Seq data were processed using *vast-tools*^103^ v2.5.1 and the VastDB release for hg38. Events with poor coverage or lacking balance between junctions were filtered (vast- tools quality column score 3 other than SOK/OK/LOW/VLOW for cassette exons, microexons, and alternative 5′ or 3′ splice site [Alt5/3] events or coverage of < 10 reads for intron retention [IR] events; score 4 other than OK/B1/B2/Bl/Bn for cassette exon and microexon events and score 5 of less than 0.05 for IR events; see https://github.com/vastgroup/vast-tools). Events with a ≥ 10% change in percent splicing between the knockdown and control samples and with a > 0.95 probability of change according to the vast-tools “diff” module were defined as differentially spliced. All RNA-Seq data and analyses have been deposited in the Gene Omnibus (GEO: GSE264355).

## Methods related to QKI functional experiments

### Plasmids

To perform transient transfections, QKI ORF, STAR domain (QKI aa 1-210 plus SV40 NLS) and LCD domain (QKI aa 211-341) were cloned into peYFP-C1 plasmid using Gibson assembly (New England Biolabs, E2611S). Mutations were introduced by site-directed mutagenesis (New England Biolabs, E0554S).

### HeLa cell culture and transfection

HeLa cells were grown in DMEM (Thermo Fisher, 11995073) with 10% FBS (Sigma) and 100 U/mL Pen-Strep (Thermo Fisher, 15140122) in a 37°C incubator supplied with 5% CO_2_ and passaged using Gibco TrypLE Express (Thermo Fisher, 12604021). For transfection, 7 × 10^4^ cells were seeded into 12-well trays (for imaging, the trays contained #1.5 thickness (0.16–0.19 mm, DIN ISO 8255 standard) coverslips). The next day, the cells in each well were transfected with 200 ng plasmid DNA, 1 µL P3000 Reagent, and 2.5 µL of Lipofectamine 3000 (Thermo Fisher). The transfection medium was replaced with fresh medium after 6 h. The cells were harvested or fixed 24 h post-transfection.

### Fluorescence *in situ* hybridisation (FISH) and immunofluorescence imaging

FISH experiments were performed as described above, but using HeLa cells instead of HEK293 cells. Following FISH, samples were washed twice with PBS and once with PBST (PBS with 0.05% Tween 20). Samples were then incubated with anti-NONO antibody (1:500, made by Monoclonal Antibody Facility / Harry Perkins Institute of Medical Research^104^) and anti-QKI antibody (1:1000, Bethyl Laboratories, A300-183A). All antibody incubations were performed for 1 hr at room temperature in the dark in a humidified chamber and all antibodies were prepared in PBST, unless otherwise stated. Samples were washed three times with PBST then incubated with Alexa Fluor 647-conjugated goat anti-mouse secondary antibody (1:300, Thermo Fisher Scientific, A-21237) and FITC-conjugated goat anti-rabbit secondary antibody (1:250, Jackson Laboratories, 111-095-006). Samples were washed three times were PBST, then incubated with PBS containing 4′,6-diamidino-2-phenylindole (DAPI; 1:15,000) at room temperature in the dark for 5min. Samples were washed once with PBS then coverslips were mounted using Vectashield antifade mounting medium (Vector Laboratories, H-1000- 10) and sealed with nail polish.

### Microscopy and image analysis

Images of NEAT1 and EYFP staining were acquired using a Delta Vision Elite Deconvolution Microscope with a 60× objective. Images were acquired as 70-step z-stacks at 0.2-μm intervals, then processed by 3D deconvolution and maximum intensity projection to generate single images for analysis. Acquisition parameters were kept consistent, as were the intensity thresholds used for samples within each experiment. Subsequent counting and quantitative fluorescent intensity analysis were performed with Nikon NiS Elements software (Version 4.3, Nikon, Tokyo, Japan).

### DRB treatment

Cells seeded on coverslips were transfected with plasmid encoding EYFP-QKI. 24 hours post transfection, cells were treated with 100 µM 5,6-Dichlorobenzimidazole 1-β-D-ribofuranoside (Sigma Aldrich, D1916) in normal growth media for 4 hours. To release transcription inhibition, DRB media was removed, cells were briefly washed twice with normal growth media and then grown normally. Timepoints were taken before DRB treatment and then 0, 20, 40, 60, 90, 120 and 240 minutes after DRB removal by fixing cells with 4% formaldehyde. Images of NEAT1, NONO and EYFP-QKI was carried out using a Delta Vision Elite Deconvolution Microscope with a 100× objective.

### RNA extraction and RT-qPCR

Cells were lysed with QIAzol Lysis Reagent (Qiagen, 79306) and RNA was extracted according to the manufacturer’s instructions. To aid the extraction of long non-coding RNAs, lysed samples were heated to 55°C and vortexed at 1,000 rpm for 10 min before adding chloroform. GlycoBlue Coprecipitant (Thermo Fisher, AM9516) was added prior to isopropanol precipitation to aid pellet visualisation. RNA samples were reverse transcribed to cDNA using the QuantiTect Reverse Transcription Kit (Qiagen, 205314). RT-qPCR reactions containing SensiMix SYBR No-ROX (Bioline, QT650-20), 250 nM of forward and reverse primers (Integrated DNA Technologies), molecular-grade H2O, and cDNA were amplified in a Rotor-Gene Q real-time PCR cycler (Qiagen). Ribosomal protein P0 (RPLP0), U6 spliceosomal RNA (U6), GAPDH, and β-actin were used as reference genes and relative mRNA expression levels were determined using the 2^−ΔΔCT^ method^105^. RT-PCR primers are listed in the extended methods.

### 3D structured illumination microscopy

SIM images were captured on a N-SIM microscope using a 100× SR Apo TIRF 1.49NA objective and an Andor iXon DU-897 EMCCD camera. Channels were captured sequentially, using 405 nm, 488 nm, and 561 nm lasers to detect DAPI, Alexa Fluor 488, and Alexa Fluor 568, respectively. Images were generated using Stack 3D SIM with five phase shifts, three angles, and z-stack increments of 0.125 µm. Images were reconstructed in NIS-Elements.

## Acknowledgements

We thank the Network Biology Collaborative Centre at the Lunenfeld-Tanenbaum Research Institute for technical assistance: B. Larsen, C. Wong, G. Liu, and J.P. Zhang for help with proteomics, and J. Tkach, M. Hasegan, and L. Brown for help with microscopy; W. Dunham, C. Go, H. Antonicka, and E. Shoubridge for generating cell lines; J. Kitaygorodsky for help with data analysis; J.R. Knight for developing visualisation tools; P. Samavarchi-Tehrani, J.P. Lambert, and G. Hesketh for method optimisation; High-Fidelity Science Communications for manuscript editing; and Lily Hansen-Gillis for help with figure design.

## Contributions

Conceptualisation, A.-C.G., B.J.A.D., B.J.B., and A.H.F.; Software, V.K., J.F.R., and B.J.A.D; Formal Analysis, B.J.A.D., V.K., S.K., M.W., and J.F.R.; Investigation, B.J.A.D, S.K., B.R.A., M.W., and Z.-Y.L.; Resources, J.-Y.Y. and C.T.; Data Curation, B.J.A.D.; Writing – Original Draft, B.J.A.D. and J.F.R.; Writing – Review & Editing, A.-C.G., with input from other authors; Visualisation, B.J.A.D., J.F.R., S.K., M.W., V.K., and B.R.A.; Supervision, A.-C.G., A.H.F., B.J.B., T.F.D., and K.R.C.; Project Administration, A.-C.G.; Funding Acquisition, A.-C.G., A.H.F., and B.J.B.

