## Supplementary figures and images for "Spatial proteomic mapping of human nuclear bodies reveals new functional insights into RNA regulation"

### Extended Data Figure 1

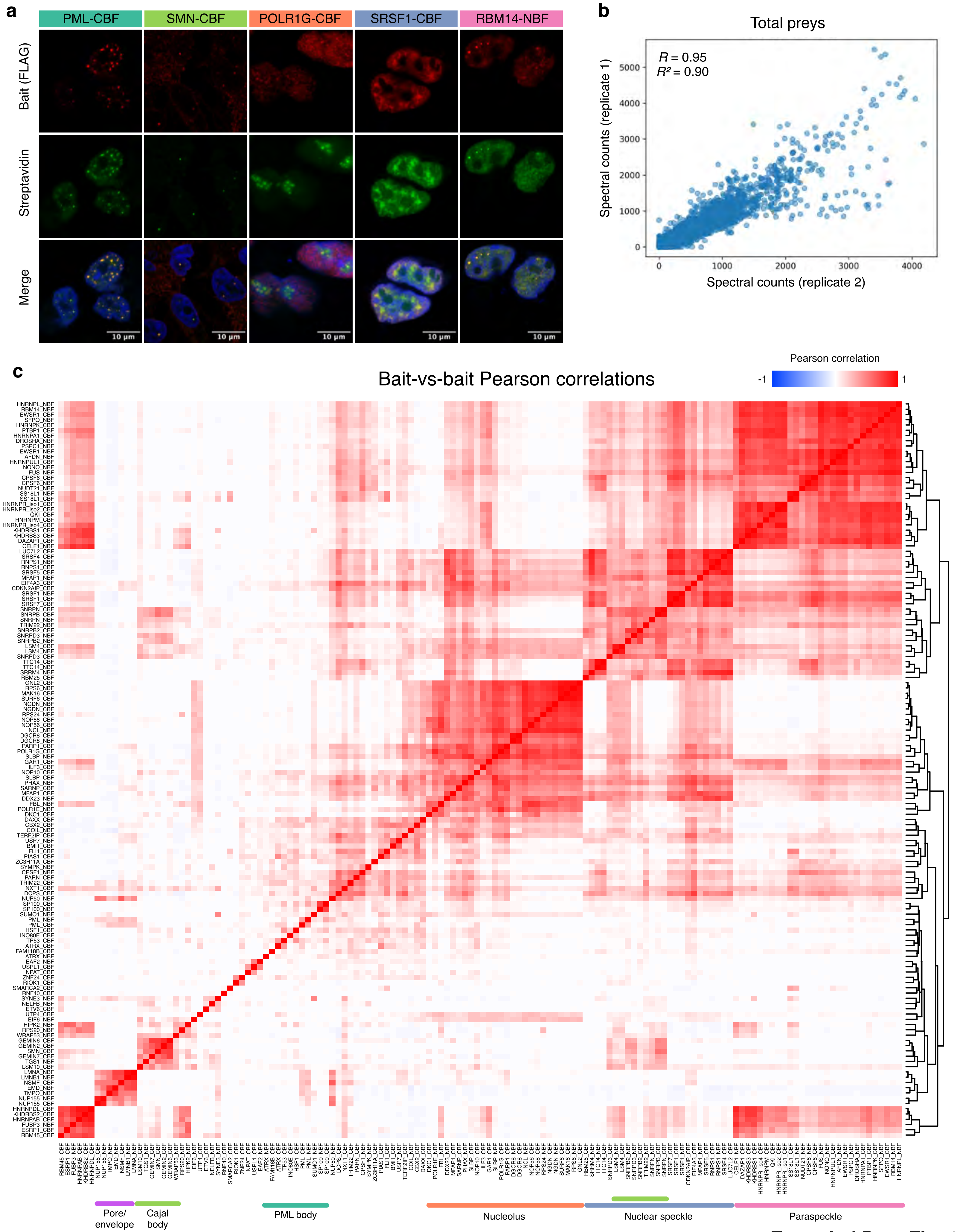


Extended Data Fig. 1

### Extended Data Figure 2

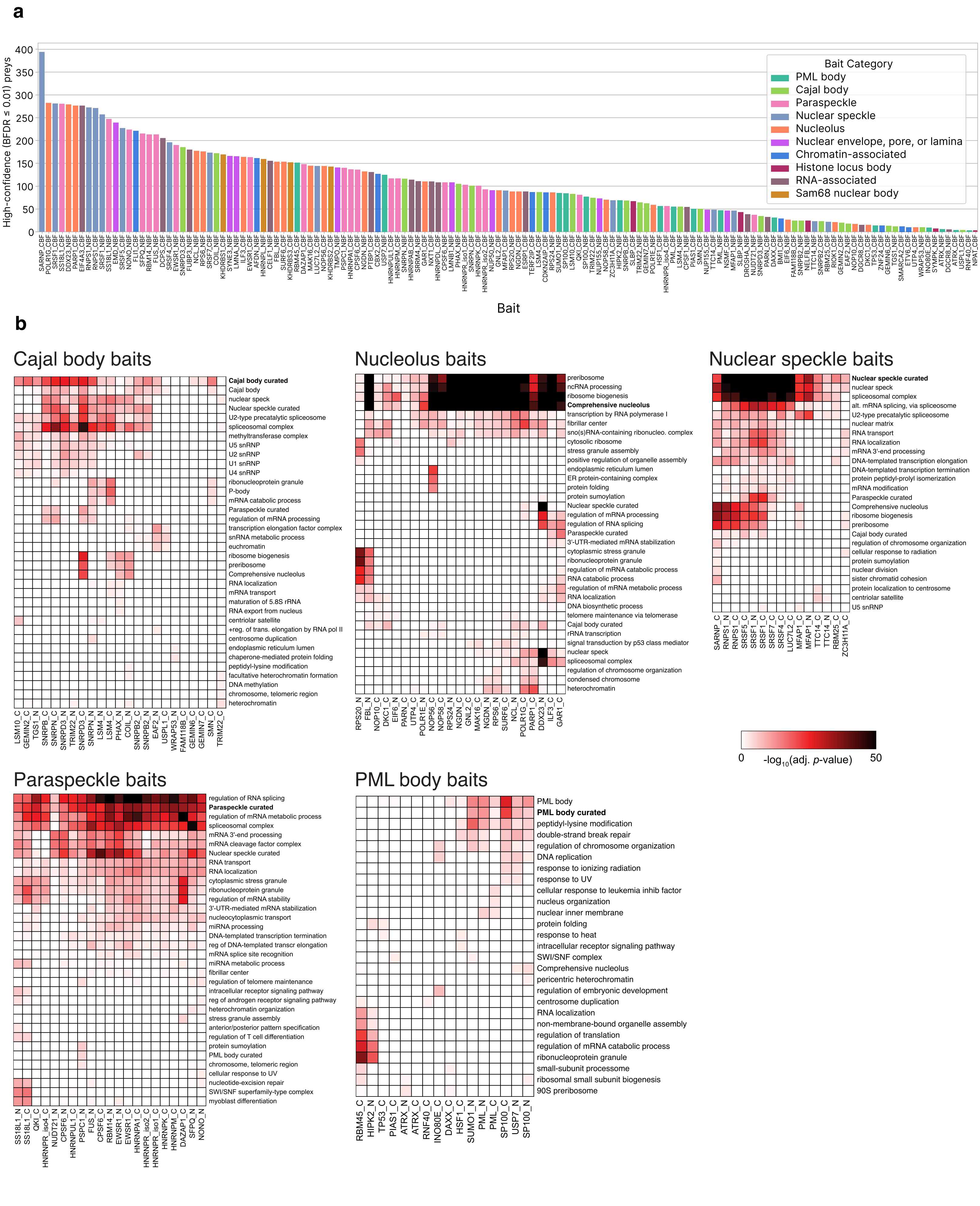


**Extended Data Fig. 2**

### Extended Data Figure 3

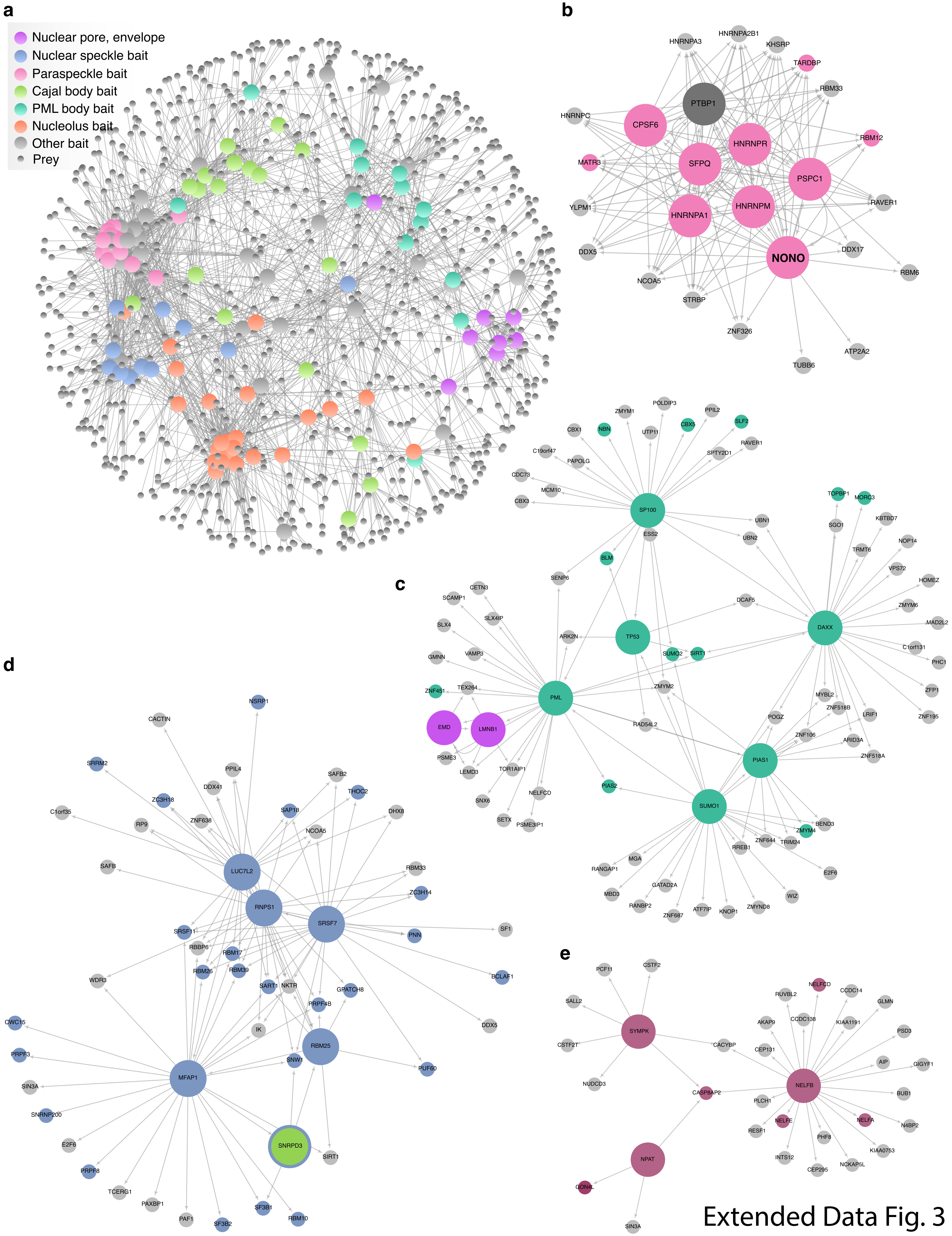


Extended Data Fig. 3

### Extended Data Figure 4

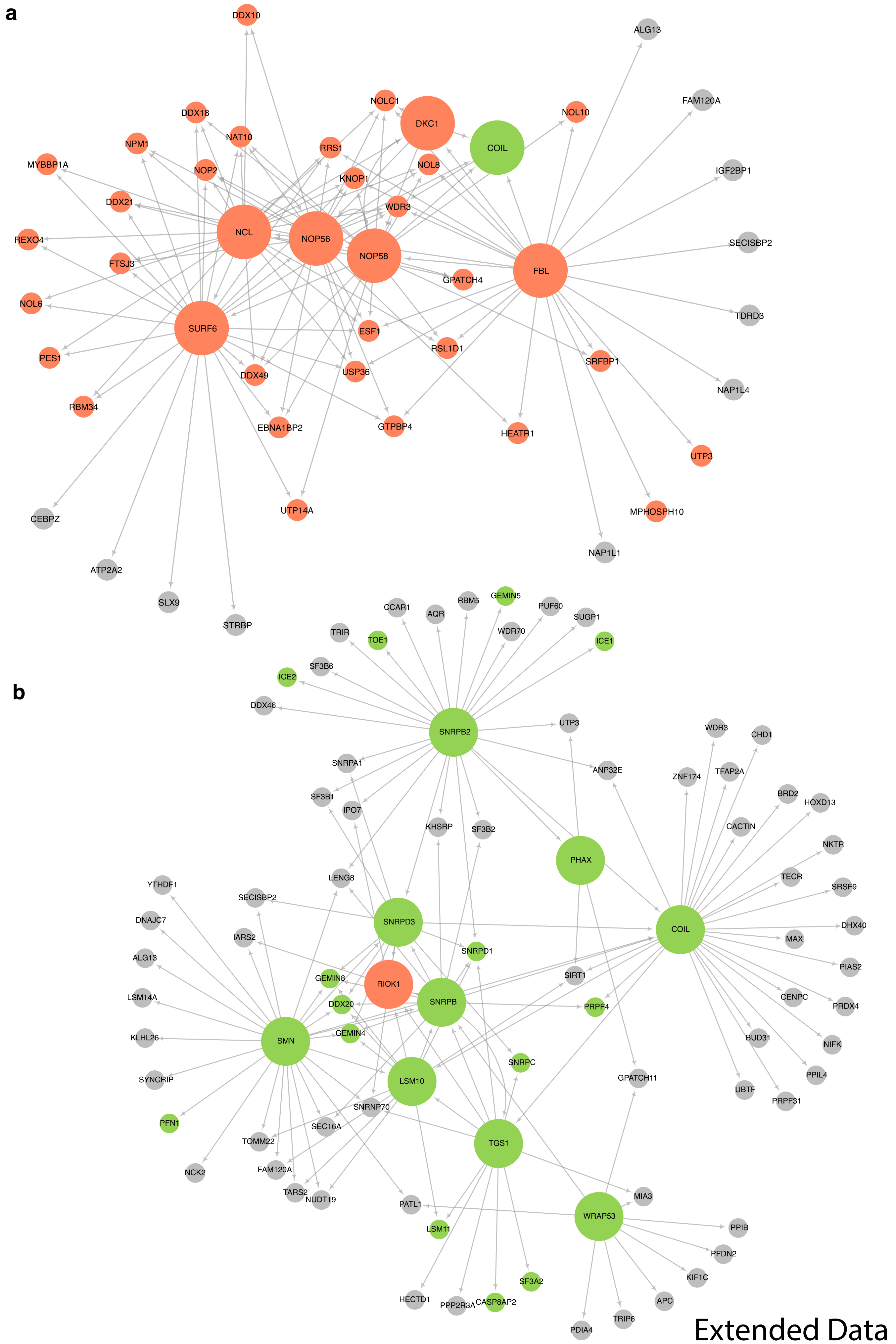


Extended Data Fig. 4

### Extended Data Figure 5

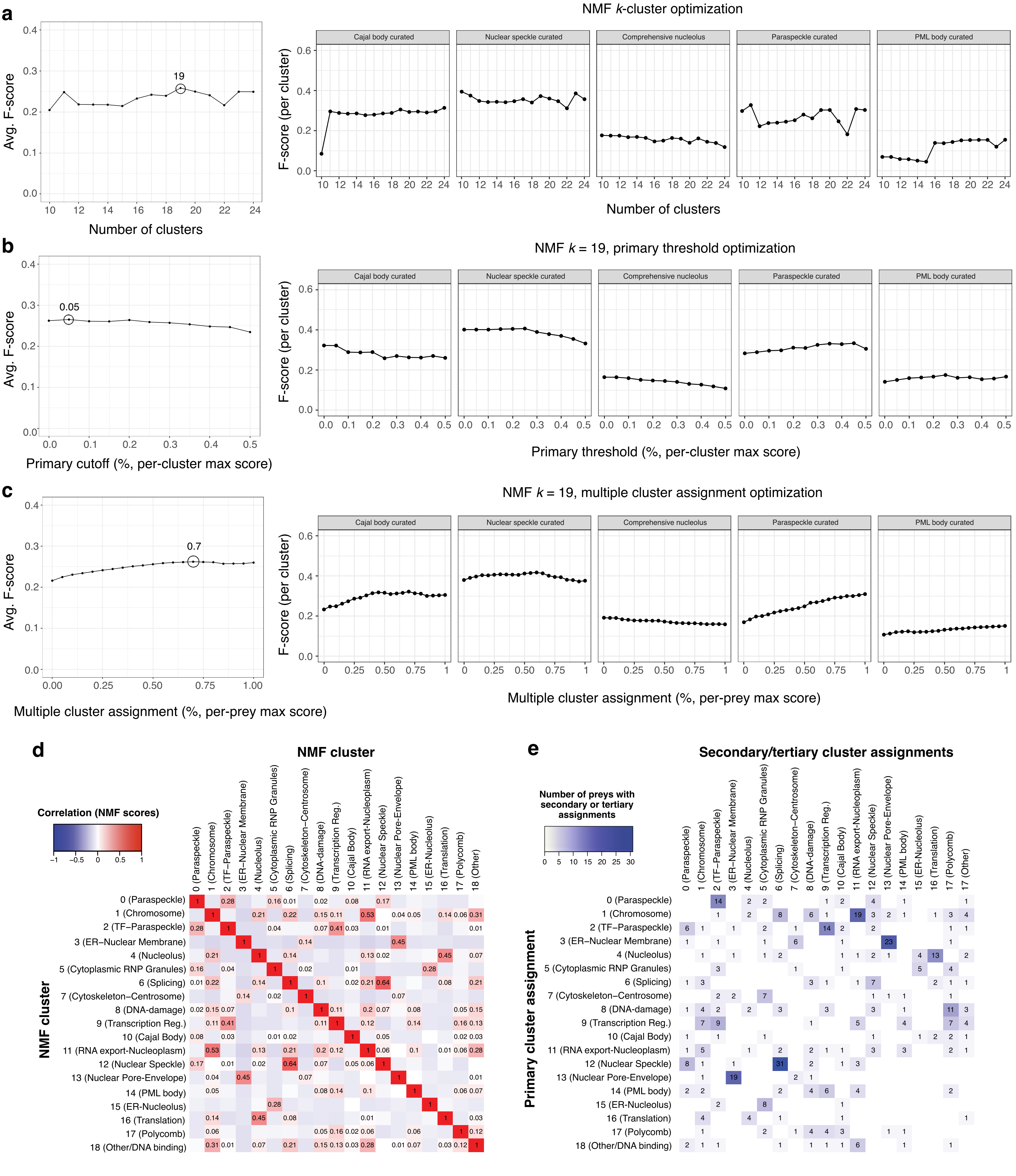


Extended Data Fig. 5

### Extended Data Figure 6

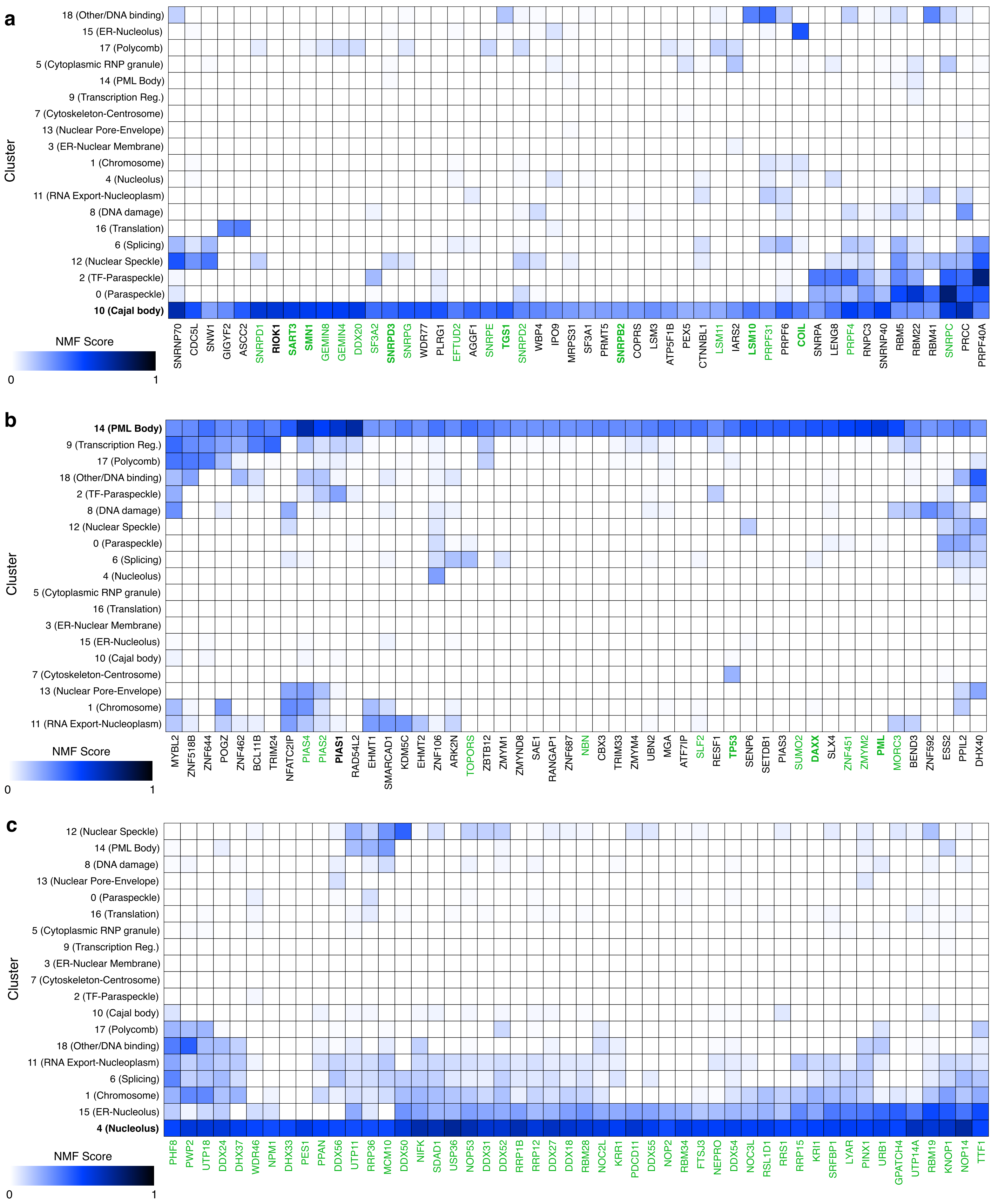


Extended Data Fig. 6

### Extended Data Figure 7

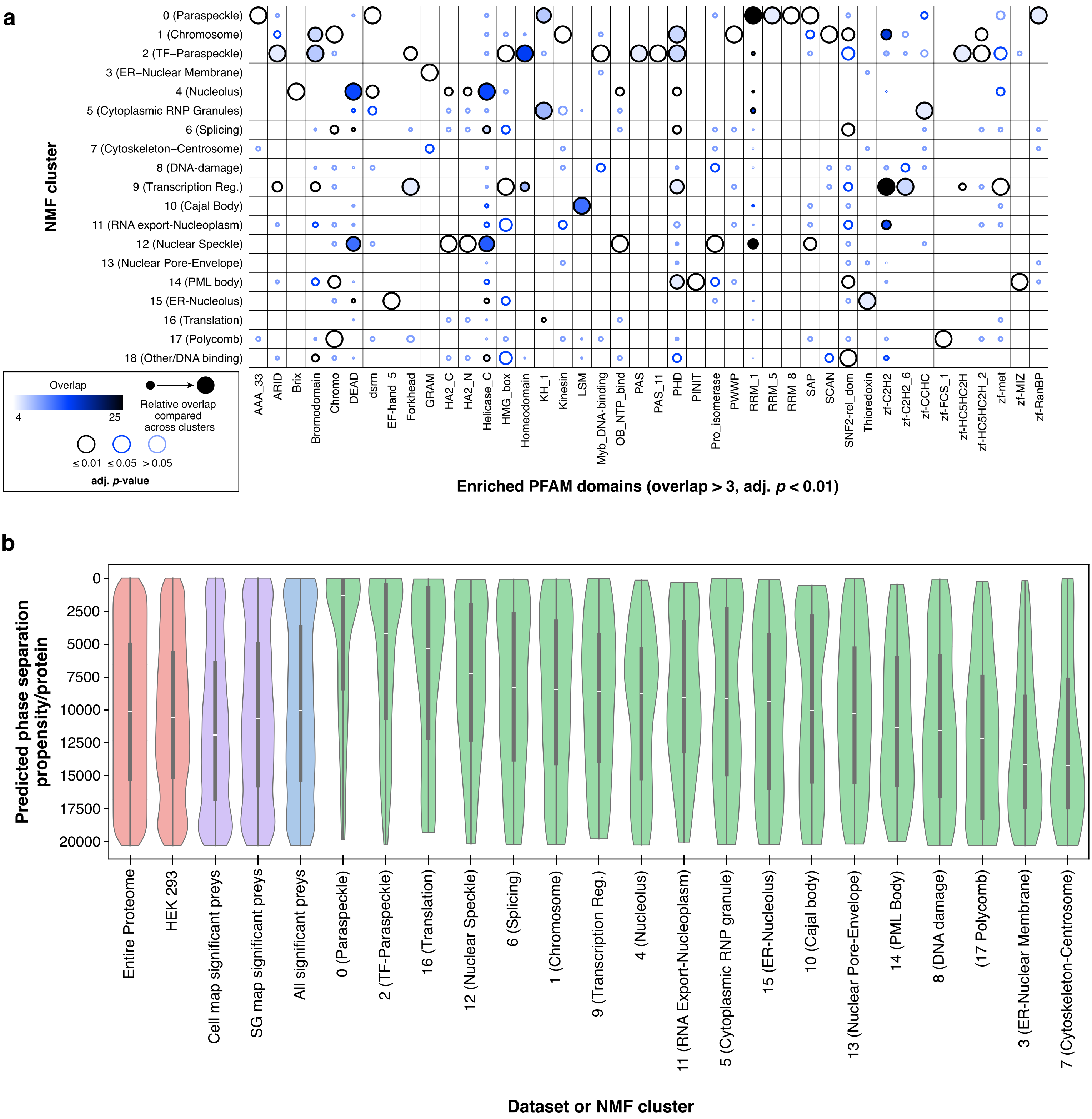


Extended Data Fig. 7

### Extended Data Figure 8

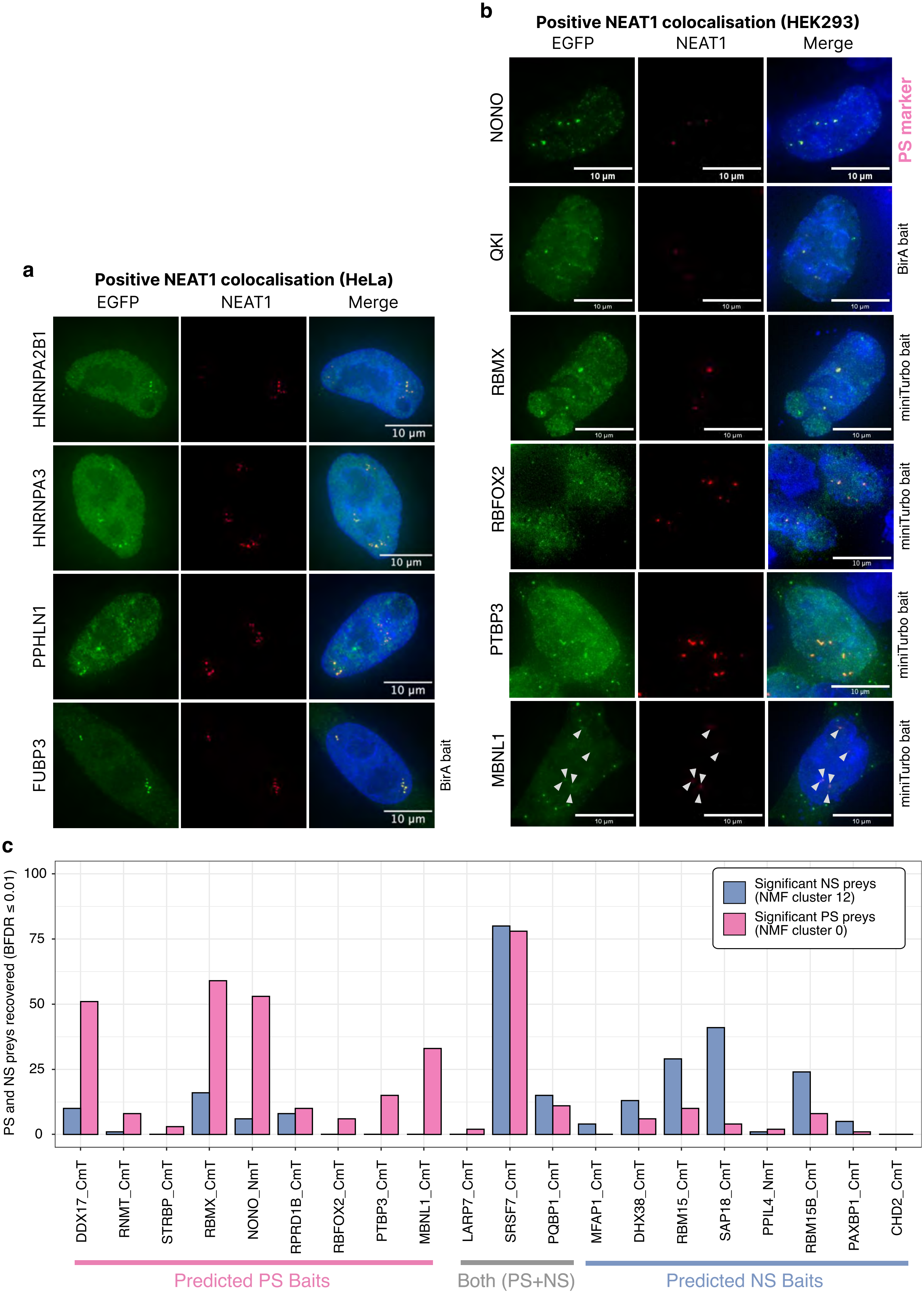


**Extended Data Fig. 8**

### Extended Data Figure 9

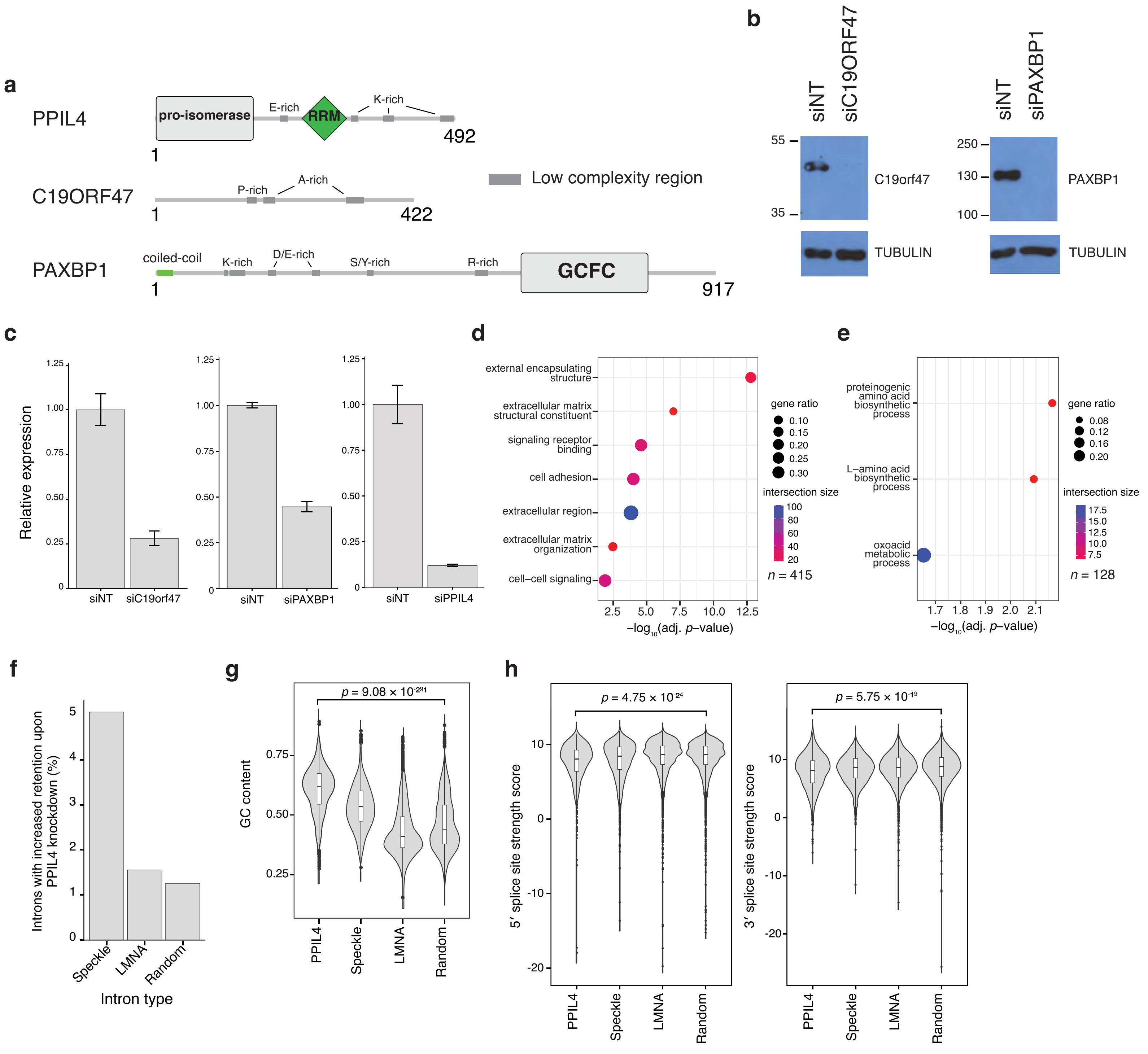


**Extended Data Fig. 9**

### Extended Data Figure 10

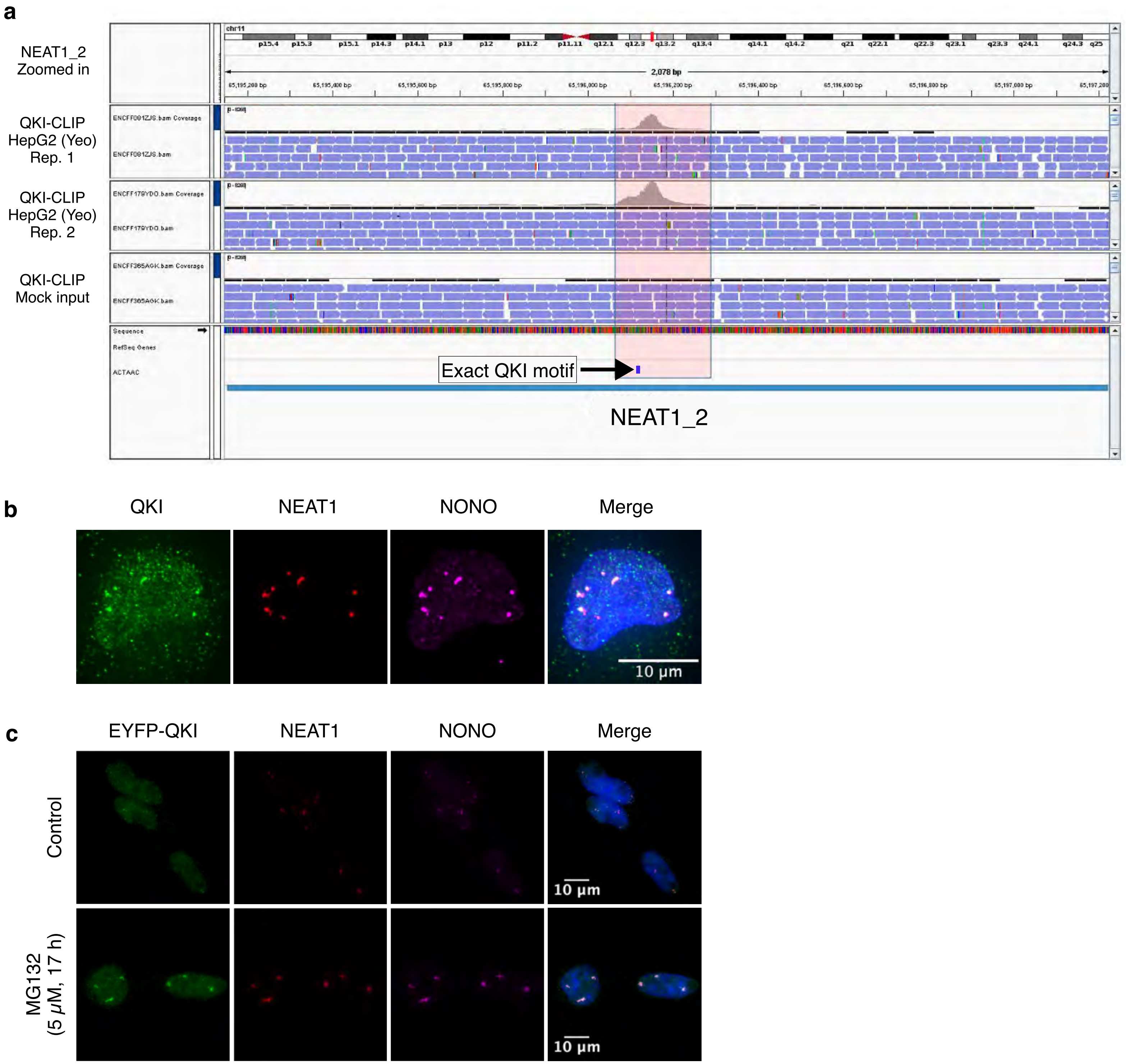


Extended Data Fig. 10
